# Synaptic Gpr85 influences cerebellar granule cells electrical properties and light-induced behavior

**DOI:** 10.1101/2025.03.10.642364

**Authors:** Romain Darche-Gabinaud, Abeer Kaafarani, Marine Chazalon, Valérie Suain, Erika Hendrickx, Louise Conrard, Anne Lefort, Frédérick Libert, Mehmet Can Demirler, Serge N. Schiffmann, David Perez-Morga, Valérie Wittamer, Marc Parmentier, Isabelle Pirson

## Abstract

GPR85/SREB2 is an exceptionally conserved orphan seven-transmembrane receptor with poorly understood biological function. Here, we combine genetic, imaging, transcriptomic, electrophysiological, and behavioral approaches in zebrafish to uncover the properties and roles of Gpr85 across development and adulthood. We show that, as in mammals, *gpr85* is expressed in diverse neuronal populations within the central nervous system, retina and intestine. Using a fluorochrome-tagged Gpr85 construct expressed in native domains, we provide the first *in vivo* evidence that Gpr85 is enriched at synaptic sites in both the brain and retina. Transcriptomic profiling of cerebellar granule cells lacking Gpr85 reveals gene expression changes consistent with increased neuronal activity. Electrophysiological recordings from cerebellar slices confirm that Gpr85-deficient granule cells exhibit heightened excitability. Functionally, Gpr85 loss enhances light-triggered startle responses in larval zebrafish. Together, these findings position Gpr85 as a synaptically enriched modulator of neuronal excitability and sensory-driven behavior, offering new insight into its roles.

## Introduction

GPR85 is classified as a class A G protein-coupled receptor (GPCR). It belongs to the Super Conserved Receptors Expressed in Brain (SREB) subfamily, which includes GPR27 (SREB1), GPR85 (SREB2) and GPR173 (SREB3). In mammals, *GPR85* shows broad expression in the brain, with particularly strong enrichment in the cerebellum (Hellebrand et al., 2000, 2001; Matsumoto et al., 2005). It is also found in the pituitary gland, retina, intestine, and testis (Ito et al., 2009; Matsumoto et al., 2000; Regard et al., 2008). Notably, *Gpr85* transcripts are detected from early developmental stages in both mouse and zebrafish (Hellebrand et al., 2001; Thisse & Thisse, 2004).

One of the most intriguing features of GPR85 is its high level of evolutionary conservation: its amino acid sequence is identical between human, primates, and rodents. Additionally, the receptor is absent in invertebrates (Hellebrand et al., 2000). In zebrafish, an evolutionarily more distant vertebrate model, the receptor nevertheless shares 93.8% amino acid identity with its human counterpart. This level of conservation strongly suggests a critical physiological role for GPR85 across vertebrates.

Despite this, GPR85 remains an orphan receptor, with no known natural ligands. In addition, no constitutive activity of the receptor has yet been conclusively demonstrated. This lack of pharmacological insight has significantly limited our understanding of GPR85’s biological functions.

From a pathological perspective, correlative studies have linked *GPR85* to psychiatric and neurodevelopmental conditions. Two *GPR85* single nucleotide polymorphisms (SNPs) have been associated with schizophrenia, thereby categorizing GPR85 as a schizophrenia risk factor (Matsumoto et al., 2008; Chen et al., 2012). Additionally, two *GPR85* missense mutations have been identified in patients with Autism Spectrum Disorder (ASD) (Fujita-Jimbo et al., 2015). Moreover, *GPR85* expression is suppressed by an edited miRNA (A to I, hsa-mir-376a-5p) that is enriched in ASD patients (Wu et al., 2022). These findings position GPR85 as a candidate modulator in the etiology of complex neurological disorders, although its mechanistic involvement remains unresolved.

Previous studies have reported that Gpr85 expression is regulated by neuronal activity. It is upregulated following treatment of neurons with excitatory (kainic acid, picrotoxin) or inhibitory (TTX) agents (Jeon et al., 2002; Jin, Kang, Kim, et al., 2018). Its expression is increased in SHANK3-overexpressing mice, an established ASD model, where its expression correlates with manic-like behavior, seizures, and altered excitation/inhibition balance (Jin, Kang, Kim, et al., 2018; Jin, Kang, Ryu, et al., 2018). These observations suggest that Gpr85 not only responds to neuronal activity but may also contribute to shaping it.

Further supporting a role at synapses, Fujita-Jimbo *et al*, (2015) reported an interaction between GPR85 and PSD-95 in cultured neurons, indicating potential postsynaptic localization. Functional studies in mouse models provided additional insights: GPR85 knock-out (KO) mice display increased brain weight and enhanced spatial memory, while mice overexpressing the receptor in postnatal forebrain neurons displayed reduced brain weight alongside social and cognitive defects (Matsumoto et al., 2008; Chen et al., 2012). These data led to the proposal that GPR85 acts as a negative regulator of adult hippocampal neurogenesis (Chen et al., 2012), although no confirmation has been reported since.

Altogether, despite promising genetic and functional data, the biological roles of GPR85 remain incompletely understood — particularly in vivo and at the systems level.

In the present work, we aimed to elucidate the endogenous properties and functions of GPR85 in a vertebrate model. To this end, we leveraged the zebrafish, a powerful system for genetic modification, live imaging, and behavior analysis, to generate Gpr85 knockout, reporter, and transgenic lines. Phenotypical characterization of these lines refined our understanding of *gpr85* expression throughout development and adulthood. Using high-resolution imaging, we revealed that Gpr85 localizes to synaptic compartments in the retina and brain. We further demonstrated that the loss of Gpr85 enhances the excitability of cerebellar granule neurons and amplifies light-induced behavioral responses.

Together, our results reveal Gpr85 as a synaptic membrane receptor that modulates neuronal activity and stimulus-evoked behavior *in vivo*, advancing our understanding of this evolutionarily conserved, yet enigmatic, receptor.

## Results

### Characterization of newly generated knock-in zebrafish *gpr85* reporter lines

Firstly, to define the tissues expressing *gpr85* in zebrafish embryos, we performed whole-mount *in situ* hybridization (WISH) at different developmental stages from 1 to 6 days post-fertilization (dpf). The signal emerged at 1 dpf in the telencephalic and ventral midbrain regions, thereafter expanding to the entire brain by 6 dpf (Appendix Fig. S1A). Transcripts were also observed in the retina of 3 dpf zebrafish larvae (Appendix Fig. S1B).

To better define *gpr85* expression, we generated a zebrafish reporter line using CRISPR/Cas9 technology with targeted integration repair (Wierson et al., 2020; Jordan M et al., 2021). The GAL4 open reading frame was inserted downstream of the endogenous *gpr85* start codon, creating a loss-of-function allele (Appendix Fig. S1C). The correct insertion of the GAL4 cassette at the *gpr85* locus was validated using PCR genotyping (Appendix Fig. S1D) and Sanger sequencing. We then introduced the *gpr85*^GAL4^ allele into either the *Tg(UAS:GFP-CAAX)^m1230^* or *Tg(UAS:LifeAct-eGFP)^mu271^* backgrounds. Phenotypic observations of these two lines did not reveal any developmental or adult behavioral defects in the heterozygous *gpr85*^GAL4/+^ state. Adult zebrafish were fertile, and their offspring followed the expected mendelian inheritance ratios. All observations described hereafter were consistent across both lines.

We first characterized the general expression profile of the GFP-CAAX reporter during the embryonic development of *gpr85*^GAL4/+^, *Tg(UAS:GFP-CAAX)* individuals. Whole body imaging of live embryos revealed that the GFP signal emerged at 1 dpf in the telencephalic and ventral midbrain regions (Fig. 1A-B). At 2 dpf, the expression expanded to include forebrain and midbrain (Fig. 1C), persisting through the end of embryogenesis (3 dpf; Fig. 1D) and into post-embryonic stages (6 dpf; Fig. 1E). Consistent with WISH experiments, GFP was detected in the retina (Fig. S1E-H’’). Immunodetection on coronal sections of 3 dpf embryos (Appendix Fig. S1E-F’’) and 6 dpf (Appendix Fig. S1G-H’’) larvae, revealed GFP-positive (GFP^+^) cells in the ganglion cell layer (GCL) and the inner nuclear layer (INL), but not in the photoreceptor layer (ONL) (Appendix Fig. S1F; H). GFP was also detected in the spinal cord (Fig. S1I-J’’), though this have not been detectable with the less sensitive WISH method (Fig. S1A-B). We observed GFP expression in both the ventral and dorsal regions of the developing spinal cord of 3 dpf embryos (Appendix Fig. S1I-I’’) and 6 dpf larvae (Appendix Fig. S1J-J’’).

**Figure 1.**
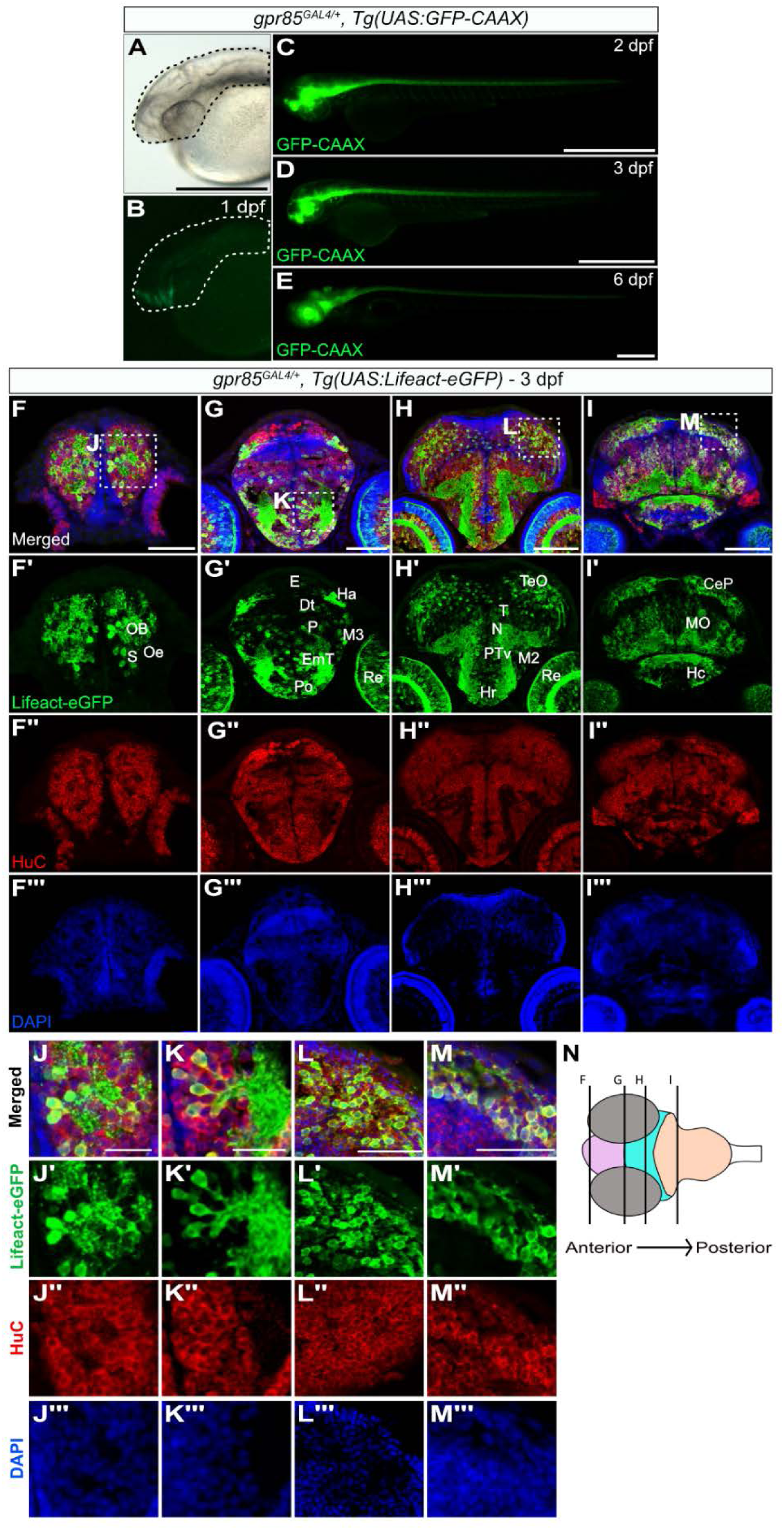
Gpr85 is broadly expressed by maturing neurons in the zebrafish brain parenchyma during late embryogenesis. A-E Lateral views of live *gpr85*^GAL4/+^, *Tg(UAS:GFP-CAAX)* zebrafish embryos at 1 dpf with brightfield (A) and fluorescence imaging (B-E) showing the GFP-CAAX signal in the brain and spinal cord at 1 dpf (B), 2 dpf (C), 3 dpf (D), and 6 dpf (E). Scale bars: 50 μm. F-I’’’ Maximum projection confocal images of coronal sections of the brain from a 3 dpf *gpr85*^GAL4/+^, *Tg(UAS:Lifeact-eGFP)* zebrafish larva stained with anti-GFP (green) (F’-I’), anti-HuC (red) (F’’-I’’) and DAPI (blue) (F’’’-I’’’). Lifeact-eGFP^+^ cells are observed in the olfactory bulb, subpallium (F’), habenula, retina, eminentia thalami, preoptic region, pallium (G’), tectum opticum, hypothalamus (H’), cerebellar plate, and medulla oblongata (I’). Boxed regions are enlarged in panels J to M. Scale Bars: 50 µm. J-M Enlarged optical sections of the areas boxed in panels (F-I), showing Lifeact-eGFP^+^/HuC^+^ neurons in the brain parenchyma. Scale bars: 25 µm. N Schematic representation of the coronal sections of the brain illustrated in F-I, with the forebrain in pink, the midbrain in turquoise, and the hindbrain in peach. Data information: CeP, cerebellar plate; DT, dorsal thalamus; EmT, eminentia thalami; H, rostral hypothalamus; Ha, habenula; Hc, caudal hypothalamus; MO, medulla oblongata; N, region of the nucleus of medial longitudinal fascicle; M2, migrated posterior tubercular area; M3, migrated area of EmT; OB, olfactory bulb; Oe, olfactory epithelium; P, pallium, Po, preoptic region; PTv, ventral part of posterior tuberculum; Re, retina; S, subpallium; TeO, tectum opticum; T, midbrain tegmentum; dpf, days post-fertilization.

Furthermore, we observed GFP signal in the intestine of 6 dpf larvae, where specific cells were labelled all along the tract (Appendix Fig. S2A-B). Using *gpr85*^GAL4/+^, *Tg(UAS:GFP-CAAX, NBT:DsRed)* fish, we identified neurons (DsRed^+^ cells) as the GFP^+^ cells in the developing intestine of 6 dpf zebrafish (Appendix Fig. S2C-D).

We then focused on the developing brain, where gpr85 was already robustly expressed by 2 dpf, to further analyze its spatiotemporal pattern.

### *Gpr85* is broadly expressed in zebrafish brain neurons at the end of embryogenesis

To delineate more precisely the anatomical structures of the developing zebrafish brain that express *gpr85*, we performed anti-GFP immunodetection on coronal sections of 3 and 6 dpf *gpr85*^GAL4/+^, *Tg(UAS:Lifeact-eGFP)* embryos. At 3 dpf, Lifeact-eGFP expression was detected throughout the brain (Fig. 1F-I). Robust expression was observed in the olfactory bulb and subpallium, while notably absent from the olfactory epithelium at this stage (Fig. 1F’). *Gpr85*-reporter expression was observed in cell clusters from left and right habenula (Fig. 1G’). In more caudal brain regions, Lifeact-eGFP^+^ cells were scattered across the brain parenchyma (Fig. 1H’-I’).

To confirm the neuronal identity of Lifeact-eGFP^+^ cells, co-immunostaining was performed with HuC, a pan-neuronal marker during zebrafish embryogenesis (Kim et al., 1996; Park et al., 2000). All Lifeact-eGFP^+^ cells were also positive for HuC (Fig. 1J-M) except for a few Lifeact-eGFP^+^/HuC^-^ cells which were observed in the midbrain ventral neuronal progenitor/precursor zone (Fig.1-supplement figure1A-A’’’, arrowheads). These cells are likely post-mitotic neuronal precursors migrating toward the developing brain parenchyma, as no PCNA^+^/Lifeact-eGFP^+^ cells were detected in this region (Fig. 1-supplement figure1B-C’’, arrowheads). Thus, at the end of embryogenesis, Lifeact-eGFP^+^ cells can largely be classified as maturing neurons.

A similar expression pattern of the *gpr85* reporter was observed in our *gpr85*^GAL4/+^*, Tg(UAS:Lifeact-eGFP)* line at 6 dpf in the hypothalamus (Fig. 1-supplement figure1D), olfactory bulb, pallium, subpallium (Fig. 1-supplement figure1E), habenula (Fig. 1-supplement figure1F; H), torus longitudinalis, optic tectum, cerebellum, and medulla oblongata (Fig. 1-supplement figure1G) (Shainer et al., 2023).

We then compared our *gpr85* expression data in embryos with a previous zebrafish brain single-cell transcriptomic dataset named “Daniocell”, which covers zebrafish development from 3 to 120 hours post fertilization (hpf) (Ja et al., 2018; Sur et al., 2023a). Consistent with our reporter line, *gpr85* transcripts were undetectable before the mid-segmentation stage (14-21 hpf) (Appendix Fig. S3A). Afterwards, transcripts were observed in the neural system, eye and spinal cord throughout embryonic and post-embryonic development (Appendix Fig. S3B). Initial expression appeared in developing forebrain neurons, motor neurons and spinal cord interneurons (Appendix Fig. S3C). At mid-segmentation, *gpr85*-expressing neurons were identified in the pallium and ventral forebrain/diencephalon (Appendix Fig. S3C).

Later during embryonic development, *gpr85* expression expanded to additional brain regions, where it was observed in various neuronal cell types, including GABAergic, glutamatergic, glycinergic and dopaminergic neurons. The expression of *gpr85* was detected in neuronal subpopulations of the retina INL and GCL, consistent with our observations (Appendix Fig. S1F and H), emerging from 36 hpf, in amacrine cells (ACs) and in ON/OFF bipolar cells (BPCs). Finally, the transcriptomic data revealed no *gpr85* expression in neural progenitors, endothelial cells, or immune cells of the central nervous system (CNS).

Taken together, these results confirm that our newly generated *gpr85*^GAL4^ reporter lines are robust and ideal tools for identifying and studying *gpr85*-expressing cell populations in the zebrafish CNS and retina.

### *Gpr85* expression is maintained in the brain, retina and intestine of adult zebrafish

The expression profile of *gpr85* in the adult zebrafish brain has been previously documented by others through qPCR analysis (Breton et al., 2023), though with limited spatial resolution. Taking advantage of our *gpr85* reporter lines, we further characterized the anatomical distribution of *gpr85*-expressing cells in the adult zebrafish brain.

We first performed tissue clearing of *gpr85*^GAL4/+^, *Tg(UAS:Lifeact-eGFP)* adult brains and analyzed sagittal and coronal sections from *gpr85*^GAL4/+^, *Tg(UAS:GFP-CAAX*) brains. As during development, GFP^+^ cells were observed in various regions of the adult brain (Fig. 2A).

**Figure 2.**
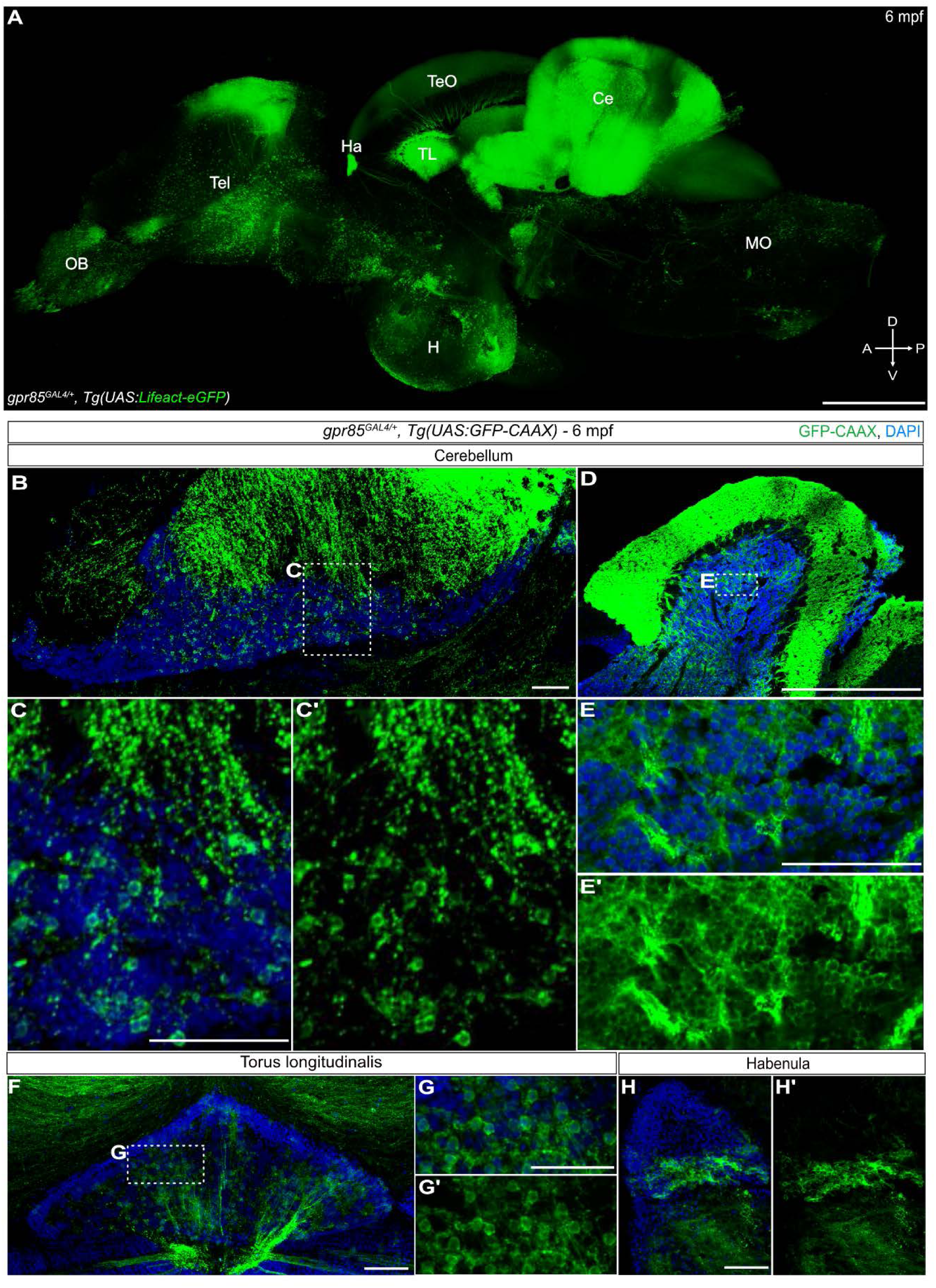
Cartography of *gpr85* expression in the brain of adult zebrafish. A Maximum projection confocal images of native Lifeact-eGFP fluorescence in a cleared adult brain hemisphere (sagittal view) from a *gpr85*^GAL4/+^, *Tg(UAS:Lifeact-eGFP)* fish. Scale bar: 500 µm. B-H’ Maximum projection confocal images of brain sections from a *gpr85*^GAL4/+^*, Tg(UAS:GFP-CAAX)* adult zebrafish stained with anti-GFP (green) and DAPI (blue). Sagittal sections of the cerebellum (B-E’) with boxes showing regions enlarged in panels C and E, respectively. Coronal section of the torus longitudinalis (F), with box highlighting the region enlarged in G. Coronal section of the habenula (H, H’). Scale bars: (B, C, E, F, G, H) 50 µm, (D) 500 µm. Data information: A, anterior; Ce, cerebellum; dpf, days post-fertilization; D, dorsal; Ha, habenula; H, hypothalamus; MO, medulla oblongata; mpf, months post-fertilization; OB, olfactory bulb; P, posterior; Tel, Telecephalum; TeO, optic tectum; TL, torus longitudinalis; V, ventral.

In the hindbrain, the highest expression was observed in the cerebellum (Ce; Fig. 2A, B-E’), with scattered GFP^+^ cells in the medulla oblongata (MO; Fig. 2A). In the midbrain, a strong signal was detected in the torus longitudinalis (TL; Fig. 2A, F-G’) and the optic tectum (TeO; Fig. 2A; Fig. 2-supplement figure1C-C’). In the forebrain, GFP^+^ cells were scattered in the olfactory bulb (OB; Fig. 2A), pallium/subpallium of the telencephalon (Tel; Fig. 2A) and the hypothalamus (H; Fig. 2A). In the hypothalamus, a strong GFP signal was observed in the posterior recess (Fig. 2A; Fig. 2-supplement figure1D-E’).

Consistent with our embryonic observations, clusters of GFP+ cells were detected in right and left habenula (Ha; Fig. 2A and H-H’). Notably, while GFP signal was prominent in the pituitary (Fig. 2-supplement figure1A-B’), GFP^+^ cells were sparse, suggesting that this signal likely arises from neuronal projections rather than from intrinsic pituitary cells.

Altogether, strong *gpr85* expression was detected in the habenula, hypothalamus, torus longitudinalis, and cerebellum of the adult zebrafish brain —mirroring patterns observed during development.

In addition to the brain, we investigated *gpr85*-reporter expression in the intestine. Using the *gpr85*^GAL4/+^, *Tg(UAS:GFP-CAAX, NBT:DsRed)* line, we confirmed the presence of GFP^+^/DsRed^+^ neurons in the adult intestine (Appendix Fig. S2E). In addition, sparse GFP^low^/DsRed^-^ cells were scattered within the intestinal villi (Appendix Fig. S2F-G).

Since Gpr85 expression has been previously reported in the human testis (Matsumoto et al., 2000), we investigated its presence in zebrafish gonads. In the testis, GFP signal was detected in all cells (Appendix Fig. S2H-H’), whereas no GFP^+^ cells were observed in the ovaries.

Consistent with embryonic findings, GFP^+^ cells were also detected in the INL and the GCL of the adult retina (Fig. 3A-A’).

**Figure 3.**
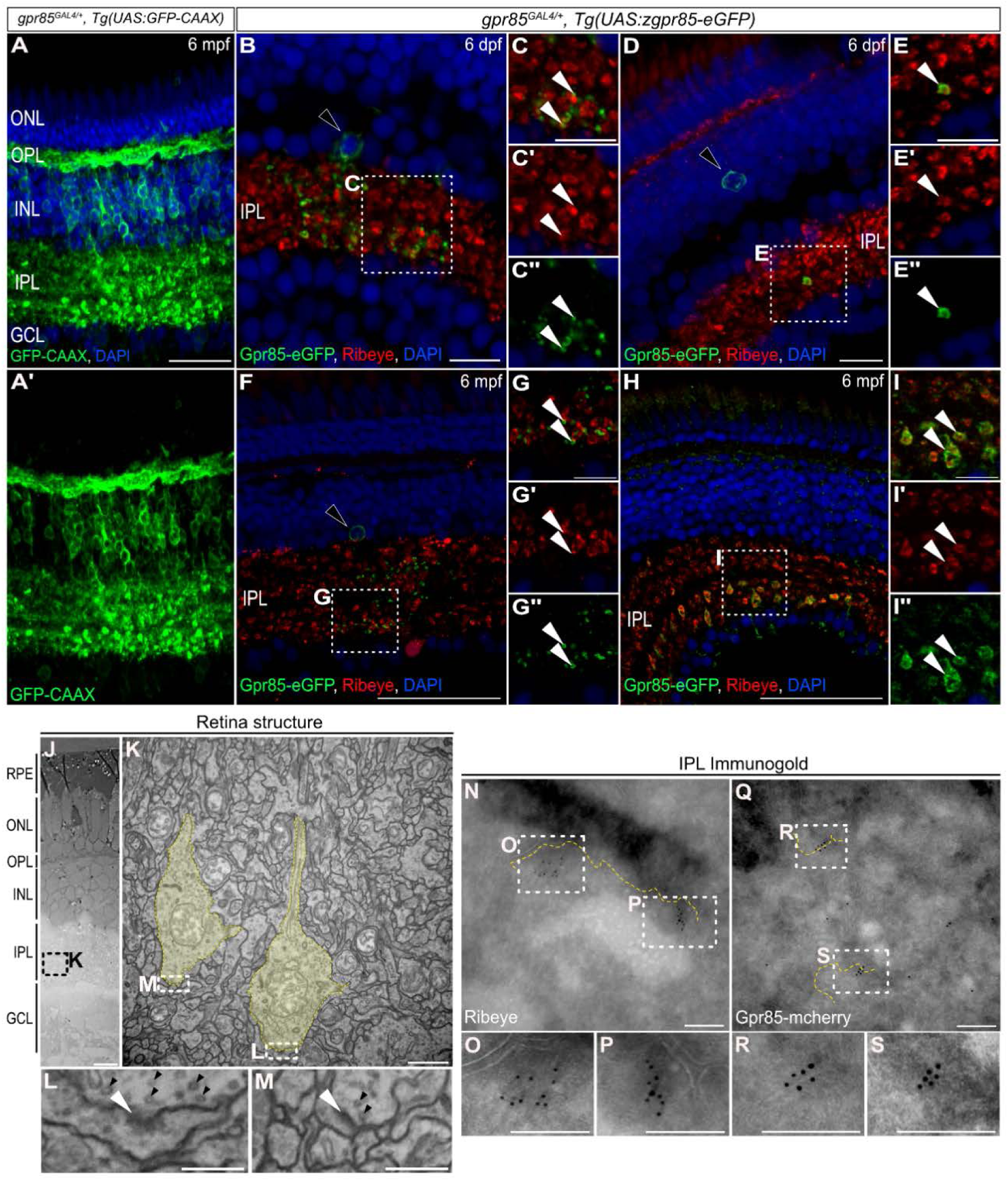
Gpr85 is enriched in the pre- and postsynaptic compartments of developing and adult IPL retinal ribbon synapses. A Maximum projection confocal images of the retina from a *gpr85*^GAL4/+^, *Tg(UAS:GFP-CAAX)* adult zebrafish stained with anti-GFP (green) and DAPI (blue). Scale bar: 50 µm. B-I’’ Confocal images of coronal retinal sections from 6 dpf (B-E’’) or 6 mpf (F-H’’) *gpr85*^GAL4/+^, *Tg(UAS:zGpr85-eGFP)* zebrafish stained with anti-GFP (green), anti-Ribeye-a (red), and DAPI (blue). (B, F) Larval or adult zGpr85-eGFP^+^ amacrine cell somas and signal within the IPL are shown, with boxed regions enlarged in C-C’’ and G-G’’, respectively. White arrowheads highlight zGpr85-eGFP signals adjacent to Ribeye^+^ ribbon terminals. (D, H) Larval or adult zGpr85-eGFP^+^/Ribeye-a^+^ bipolar cell ribbon pre-synaptic terminals are shown, with boxed regions enlarged in E-E’’ and I-I’’, respectively. White arrowheads show examples of zGpr85-eGFP signal present in Ribeye^+^ ribbon terminals. Scale bars: (B,D) 25 µm, (F, H) 50 µm, (C, E, G, I) 10 µm. J Electron microscopy coronal view of the structure of 6 dpf retina from *gpr85*^GAL4/+^, *Tg(UAS:zGpr85-mCherry)* zebrafish larvae. Scale bar: 10 µm. K Enlarged image of the ribbon pre-synaptic terminals (yellow) from the boxed area in panel J. Boxes indicate regions enlarged in panels M and L. Scale bar: 1 µm. L-M Enlarged images of synaptic boutons from the boxed area in panel K. White arrowheads highlight postsynaptic densities while black arrowheads point to presynaptic vesicles. Scale bars: 200 nm. N-Q Immunogold labeling of Ribeye (N-P) and mCherry-tagged Gpr85 (Q-S) focusing on the retinal inner plexiform layer from 6 dpf *gpr85*^GAL4/+^, *Tg(UAS:zGpr85-mCherry)* larvae. Dashed yellow lines highlight plasma membranes. Boxes indicate regions enlarged in panels O, P, R, and S. Scale bars: 200 nm. O-P Enlarged images of the boxed areas in panel N. Ribeye is successfully detected in the IPL and appears clustered near the membrane of bipolar cell terminals. Scale bars: 200 nm. R-S Enlarged images of the boxed areas in panel Q. mCherry-tagged Gpr85 appears clustered near plasma membranes in the IPL. Scale bars: 200 nm. Data information: dpf, days post fertilization; mpf, months post fertilization; ONL, outer nuclear layer; OPL, outer plexiform layer; INL, inner nuclear layer; GCL ganglion cell layer.

Taken together, these results show *gpr85* expression is maintained from late embryogenesis through adulthood in the brain, retina, and intestine, supporting its persistent and region-specific role in the zebrafish nervous system and gut.

### Gpr85 receptor localizes to synapses in the retina and brain

To gain insight into the biological function of Gpr85, we characterized its subcellular localization *in vivo*, with a focus on neurons of the zebrafish brain and retina. A previous *in vitro* study suggested an interaction between GPR85 and neuroligin-bound PSD-95, a postsynaptic marker of excitatory synapses (Fujita-Jimbo et al., 2015). Based on this, we hypothesized that Gpr85 localizes to chemical synapses in zebrafish neurons.

To test this, we used our *gpr85*^GAL4/+^ line to overexpress the Gpr85 receptor fused to either eGFP or mCherry at its C-terminal end. Both *gpr85*^GAL4/+^, *Tg(UAS:zGpr85-eGFP)* and *gpr85*^GAL4/+^, *Tg(UAS:zGpr85-mCherry)* lines were fertile and did not exhibit developmental defects. While no ectopic expression was observed, both lines exhibited mosaic labeling compared to the Lifeact-eGFP and GFP-CAAX reporter lines.

We first assessed subcellular localization in the retina. Coronal sections of the retina from *gpr85*^GAL4/+^, *Tg(UAS:zGpr85-eGFP)* fish revealed zGpr85-eGFP^+^ ACs (Fig. 3B and F) and BPCs (Fig. 3D and H) during development and adulthood. In ACs, the zGpr85-eGFP signal was localized in the soma (Fig. 3B and F, black arrowhead) and dendritic arborization of the cells within the IPL at both 6 dpf (Fig. 3B) and 6 months post fertilization (mpf) (Fig. 3F). In BPCs, the zGpr85-eGFP signal was also observed in the soma (Fig. 3D, black arrowhead) and neurites/terminals in larvae IPL (Fig. 3D) while only found in the IPL of adult zebrafish (Fig. 3H). The dendritic arborization of BPCs was essentially devoid of zGpr85-eGFP.

To assess synaptic localization, we performed co-immunostaining for αRibeye-a, a presynaptic ribbon synapse marker. zGpr85-eGFP signal was enriched at the postsynaptic sites of ribbon synapses in the dendritic arborization of developing (Fig. 3C-C’’, white arrowheads) and adult (Fig. 3G-G’’, white arrowheads) ACs. In developing (Fig. 3E-E’’, white arrowheads) and adult (Fig. 3I-I’’, white arrowheads) BPCs, zGpr85-eGFP displayed a strong signal at the level of the Ribeye-a^+^ presynaptic terminals of ribbon synapses.

These findings provide the first *in vivo* evidence that Gpr85 is enriched at both pre- and postsynaptic sites of ribbon synapses in the developing and adult retina.

Given that Gpr85 is a seven-transmembrane receptor, it was anticipated to localize at the plasma membrane *in vivo*. However, no direct evidence of such localization had been reported previously. Using *gpr85*^GAL4/+^, *Tg(UAS:zGpr85-mCherry)* larvae, we investigated the distribution of the receptor by electron microscopy using immunogold staining. The initial characterization of the retinal structure revealed the ribbon terminals within the IPL (Fig. 3J-K, yellow). Presynaptic compartments were identifiable due to the presence of presynaptic vesicles (Fig. 3L-M, black arrows) while postsynaptic densities were identified as thick and electron-dense structures (Fig. 3L-M, white arrowheads). Ribeye-a immunogold labeling confirmed presynaptic identity (Fig. 3N-P). zGpr85-mCherry exhibited localization at the plasma membrane in the IPL (Fig. 3Q-S) confirming its membrane localization *in vivo*.

Next, we assessed the subcellular localization of the chimeric receptor in the developing and adult brain, using the *Tg(UAS:FinGR(PSD)-GFP)* and *Tg(UAS:FinGR(GPHN)-mCherry)* reporter lines to discriminate between excitatory and inhibitory synapses, respectively (Son et al., 2016). These lines were crossed to *gpr85*^GAL4/+^, *Tg(UAS:zGpr85-mCherry)* or *Tg(UAS:zGpr85-eGFP)* backgrounds. Co-immunostaining was performed on 6 dpf coronal sections and adult sagittal sections of the brain.

As in the retina, zGpr85-eGFP^+^ and zGpr85-mCherry^+^ puncta were identified in neuronal projections at both larval and adult stages (Fig. 4). We observed both zGpr85-mCherry^+^/PSD^+^ (Fig. 4A,B, white arrowheads) and zGpr85-eGFP^+^/GPHN^+^ (Fig. 4C,D, white arrowheads) synapses in multiple regions of the developing and adult brain, including the telencephalon, hypothalamus, optic tectum, and cerebellum.

**Figure 4.**
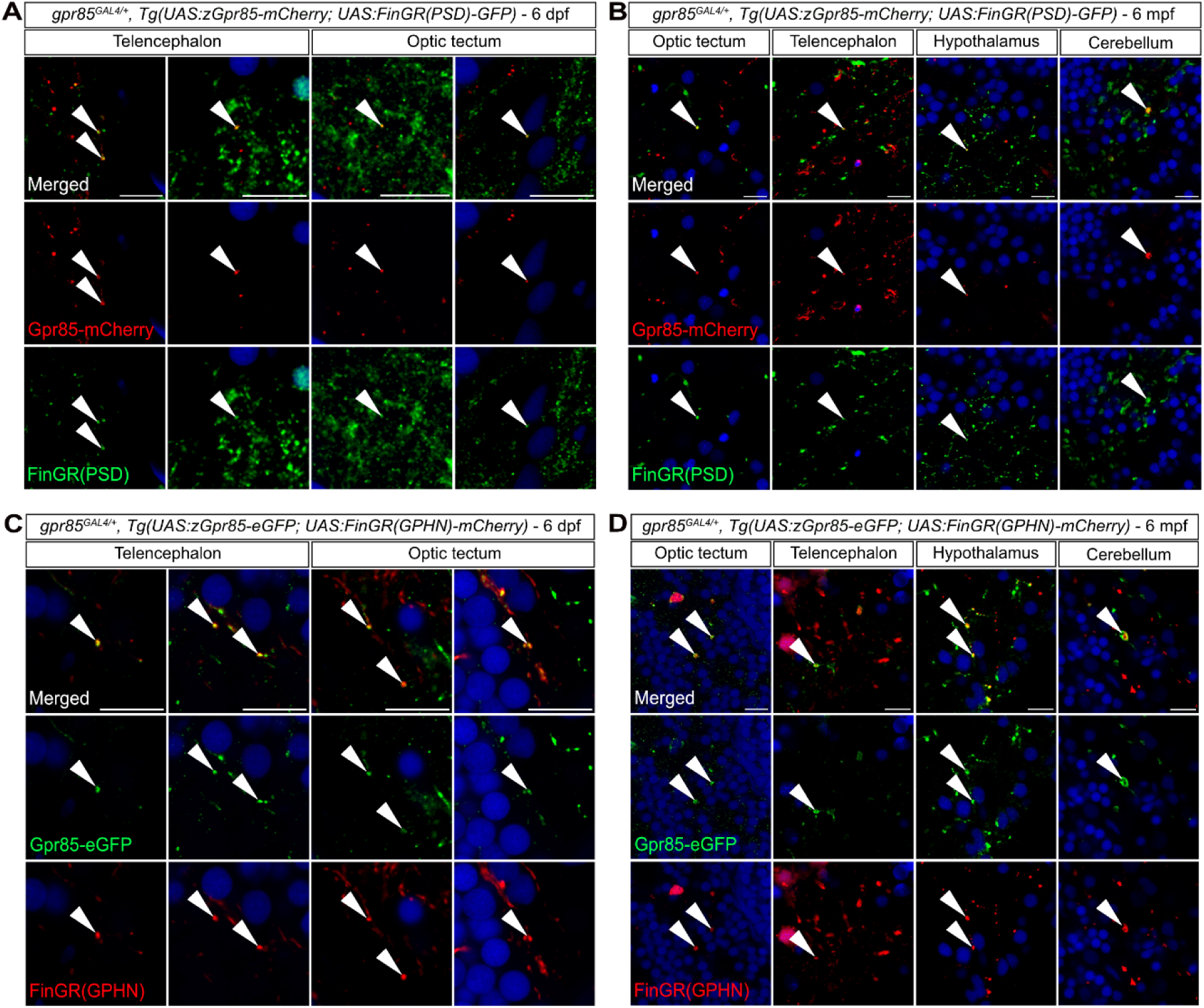
Gpr85 is enriched at the level of excitatory and inhibitory synaptic inputs of *gpr85*-expressing neurons throughout the developing and adult brain. A Confocal images of coronal brain sections from 6 dpf *gpr85*^GAL4/+^, *Tg(UAS:zGpr85-mCherry, UAS:FinGR(PSD)-GFP)* larvae stained with anti-GFP (green), anti-mCherry (red), and DAPI (blue). White arrowheads show examples of zGpr85-mCherry signal localized at the level of PSD^+^ excitatory synapses in the telencephalon and optic tectum. Scale bar: 10 µm. B Confocal images of coronal brain sections from 6 mpf *gpr85*^GAL4/+^, *Tg(UAS:zGpr85-mCherry, UAS:FinGR(PSD)-GFP)* adult zebrafish stained with anti-GFP (green), anti-mCherry (red) and DAPI (blue). White arrowheads show examples of zGpr85-mCherry signal localized at the level of PSD^+^ excitatory synapses in the indicated brain regions. Scale bar: 10 µm. C Confocal images of coronal brain sections from 6 dpf *gpr85*^GAL4/+^, *Tg(UAS:zGpr85-eGFP, UAS:FinGR(GPHN)-mCherry)* larvae stained with anti-GFP (green), anti-mCherry (red), and DAPI (blue). White arrowheads highlight examples of zGpr85-eGFP signal present at the level of the GPHN^+^ inhibitory synapses in the telencephalon and optic tectum. Scale bar: 10 µm. D Confocal images of coronal brain sections from 6 mpf *gpr85*^GAL4/+^, *Tg(UAS:zGpr85-eGFP, UAS:FinGR(GPHN)-mCherry)* adult zebrafish stained with anti-GFP (green), anti-mCherry (red) and DAPI (blue). White arrowheads highlight examples of zGpr85-eGFP signal present at the level of the GPHN^+^ inhibitory synapses of the indicated brain regions. Scale bar: 10 µm. Data information: dpf, days post fertilization; mpf, months post fertilization.

Overall, our findings provide the first *in vivo* evidence that Gpr85 is a synaptic membrane receptor, localized at both excitatory and inhibitory chemical synapses in the zebrafish retina and CNS across development and adulthood.

### Gpr85 is dispensable for the development, viability, and formation of excitatory inputs of *gpr85*-expressing optic tectal neurons

We previously showed that *gpr85* is expressed in multiple differentiating neuronal subpopulations during early zebrafish brain development. In addition, we provided evidence that Gpr85 is a synaptic membrane receptor *in vivo*. These findings raised the question of whether Gpr85 is required for the development or maturation of *gpr85*-expressing neurons.

To address this, we generated a complementary constitutive Gpr85-KO model (*gpr85*^Δ4/Δ4^) (Appendix Fig. S4A). Similar to the *gpr85*^GAL4/GAL4^ line, homozygous mutants were viable, fertile, and displayed no noticeable developmental defects (Appendix Fig. S4B,C). The offspring of these mutants followed a Mendelian inheritance ratio (Appendix Fig. S4D).

Although a prior study reported increased brain weight in GPR85-KO mice (Matsumoto et al., 2008), we found no significant differences in body length or in the wet brain weight-to-body length ratio in either of the Gpr85-KO zebrafish models (Appendix Fig. S4E,F). Nevertheless, we observed that *gpr85* transcript levels were approximately twice as high in both Gpr85-KO models compared to WT 6 dpf larvae (Appendix Fig. S4G), suggesting that the absence of functional Gpr85 triggers upregulation of the gene and/or increases the stability of its transcripts.

To further investigate the impact of Gpr85-KO on the development of *gpr85*-expressing neurons, we used *gpr85*^GAL4/Δ4^, *Tg(UAS:FinGR(PSD)-GFP)* larvae to quantify the developing optic tectal *gpr85*-expressing neurons in the *stratum paraventriculare* (SPV) and their excitatory inputs within the neuropil. The neuropil, almost devoid of neuronal soma, exhibits a uniform distribution of excitatory inputs at 6 dpf (Appendix Fig. S4H-M).

We observed no significant changes in the density of FinGR(PSD)-GFP^+^ cells in the SPV of 6 dpf larvae in the Gpr85-KO larvae compared to controls (Appendix Fig. S4I,J). Similarly, the density of FinGR(PSD)-GFP^+^ synaptic inputs within the tectal neuropil remained unaffected in whole-mount immunostaining of Gpr85-KO larvae (Appendix Fig. S4L,M).

These results show that Gpr85 is dispensable for the development, viability and formation of excitatory inputs in *gpr85*-expressing neurons during zebrafish development.

### Transcriptomic analysis of Gpr85-KO cerebellar granule cells reveals changes in genes related to neuronal activity

Although Gpr85 appears dispensable for the development of *gpr85*-expressing neurons and developmental synaptogenesis, it may influence neuronal homeostasis and/or synaptic activity. To explore the potential impact of constitutive Gpr85 deficiency on adult *gpr85*-expressing neurons, we conducted single-cell RNA sequencing (scRNAseq) on eGFP^+^ sorted cells from the adult brains of *gpr85*^GAL4/GAL4^, *Tg(UAS:Lifeact-eGFP)* and *gpr85*^GAL4/+^, *Tg(UAS:Lifeact-eGFP)* zebrafish.

After quality control and filtering, data from both genotypes were merged for clustering, which identified 11 distinct clusters under both control and knock-out conditions (*gpr85*^GAL4/+^, Ctl, n = 4219 cells; *gpr85*^GAL4/GAL4^, KO, n = 3341 cells) (Fig. 5A). As expected, 96% of cells expressed the eGFP transgene (Fig. 5B). *Gpr85* transcripts were detected, though not in all cells (Fig. 5B), likely due to expression levels below the detection threshold of the sequencing method and/or dynamic expression patterns not captured by the stable eGFP reporter. Unlike our RT-qPCR findings at 6 dpf, this adult dataset showed no global or cluster-specific upregulation of *gpr85* transcripts in mutants.

**Figure 5.**
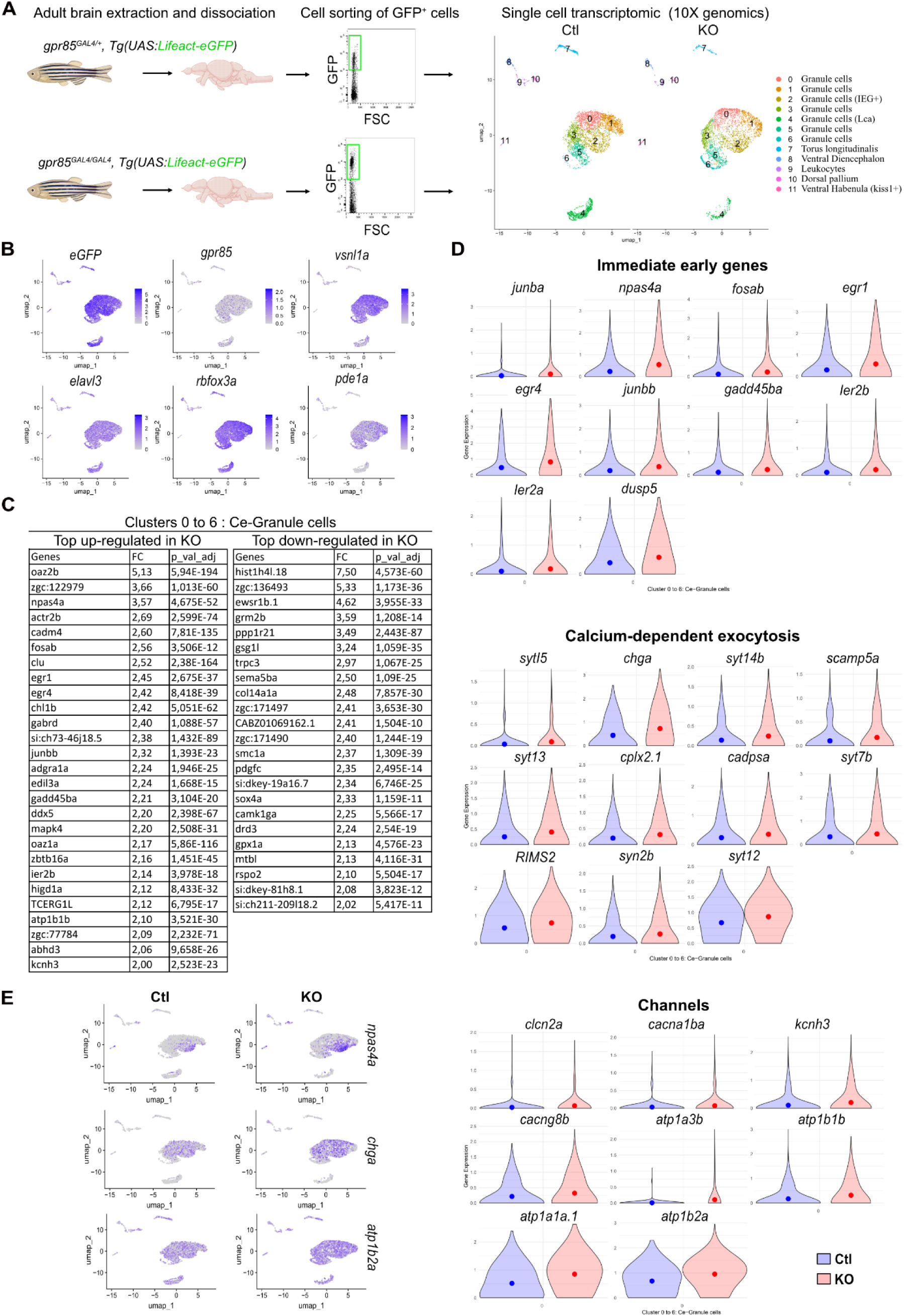
scRNAseq analysis of eGFP^+^ sorted cells from adult *gpr85*^GAL4/+^ and *gpr85*^GAL4/GAL4^, *Tg(UAS:lifeact-eGFP)* dissociated brains reveals changes in gene expression related to neuronal activity. A Experimental strategy for assessing transcriptomic changes in *gpr85*-expressing cells from the Gpr85-deficient adult brain. GFP+ cells were sorted from *gpr85*^GAL4/+^ (Ctl) and *gpr85*^GAL4/GAL4^ (KO), *Tg(UAS:lifeact-eGFP)* dissociated brains (cell sorting strategy shown with 10,000 events per condition). The right panel shows the UMAPs of cells from the Ctl and KO conditions (n = 4219 cells and n = 3341 cells, respectively) after filtering and clustering. B UMAPs of Ctl and KO cells merged, displaying expression of the UAS:lifeact-eGFP transgene, *gpr85*, the two pan-neuronal markers *elavl3* and *rbfox3a*, as well as the expression of the granule cell markers, vsnl1a and pde1a. C Table of the most up- and down-regulated genes within the isolated cerebellar clusters (0 to 6) in the KO condition, with a minimal fold change of two and an adjusted p-value < 10^-4^, expressed by at least 6% of the cells. D Violin plots of selected differentially expressed genes (DEGs) with a minimal fold change of 1.4 and an adjusted p-value < 10^-4^. All genes referenced are upregulated (mean represented by the dots). Upper panel: genes documented as immediate early genes. Intermediate panel: selection of genes related to calcium-dependent exocytosis. Lower panel: selection of genes encoding channels related to neuronal excitability/activity (voltage-dependent channels: *clcn2a*, *cacna1ba*, *kcnh3*, *cacng8b*; Na^+^/K^+^ ATPase subunits: *atp1a3b*, *atp1b1b*, *atp1a1a.1*, *atp1b2a*, *syn2b*). E UMAPs of *npas4a*, *chga* and *atp1b2a* expression in Ctl and KO conditions, respectively.

All clusters, except cluster 9, displayed neuronal identities, expressing the pan-neuronal markers *elavl3* and *rbfox3a* (Fig. 5B). Based on *gad1a* expression, we observed that 98% of the cells were non-GABAergic neurons. Cluster 9 was identified as non-neuronal, expressing leukocyte and microglial markers such as *cd74b*, *mhc2a*, *apoeb* (Fig. 5-supplement figure1 A, B).

Clusters 0–6 were identified as cerebellar granule cells (GCs) based on markers like *vsnl1a* and *pde1a* (Fig. 5B, Fig. 5-supplement figure1 A,B), while lacking markers for Purkinje, eurydendroid, Golgi and stellate cells (*gad1a, calb2a, aldoca, ca8, slc1a3b, fabp7a*) (Fig. 5-supplement figure1 C). Cluster 8 was composed of ventral telencephalon neurons, cluster 10 of dorsal pallium neurons characterized by expression of *eomesa* (Fig. 5-supplement figure1 B), and cluster 11 of *kiss1*+ neurons from the ventral habenula (Fig. 5-supplement figure1 B). Overall, ∼90% cells were classified as cerebellar GCs.

To identify transcriptional changes resulting from the loss of Gpr85, we focused on cerebellar GCs from clusters 0 to 6 to assess differential expressed genes (DEGs). Firstly, we selected genes with a fold change (FC) ≥ 2, adjusted p-value < 10^-4^ and expressed by ≥ 6% of cells. We identified 27 upregulated and 23 downregulated genes (Fig. 5C). The most upregulated gene was the ornithine decarboxylase antizyme *oaz2b* (FC = 5.13), and *oaz1a,* was also among the upregulated genes. The histone-like *hist1h4l.18* was the most downregulated gene.

Remarkably, seven of the most upregulated DEGs were identified as immediate early genes (IEGs): *npas4a*, *fosab*, *egr1*, *egr4*, *junbb*, *gadd45ba* and *ier2b* (Fig. 5C). Basal expression of *fosab* and *egr1* transcription factors in the adult zebrafish cerebellum has been reported (Kühn and Köster, 2010; Kress and Wullimann, 2012). Consistent with this, several of these IEGs were defined as main markers specifying cluster 2, confirming the existence of a GC subpopulation expressing multiple IEGs in basal conditions (Fig. 5D, Fig 5-supplement figure1 A, B, Appendix Table S1).

Additional upregulated genes included three genes encoding proteins involved in neuronal activity and excitability, a potassium voltage-gated channel (*kcnh3*), an ATPase Na^+^/K^+^ transporting subunit (*atp1b1b)* and the gamma-aminobutyric acid type A receptor subunit delta (*gabrd*) (Fig. 5C). Based on these findings, we examined the DEG list for further enrichment in genes related to neuronal activity and excitability.

In total, we observed ten documented IEGs which displayed a significant upregulation in the Gpr85-KO GCs, mostly observed within the IEGs^+^ GCs cluster 2 (Fig. 5D, E, Appendix Table S1, FC > 1.7, adjusted p-value < 10^-11^). In addition, there was upregulation of 11 genes related to neuronal activity-dependent exocytosis, and 8 genes encoding ATPase Na^+^/K^+^ exchangers or voltage-dependent channels (Fig. 5D, Appendix Table S1, FC > 1.4, adjusted, p-value < 10^-4^). Unlike the IEGs, upregulation of the channels was broadly distributed across GCs clusters, as exemplified with *npas4a*, *chga* and *atp1b2a* (Fig. 5E).

Altogether, these findings confirm that *gpr85* is predominantly expressed in cerebellar GCs within the adult zebrafish brain. The absence of Gpr85 alters the transcriptional profile of these neurons, particularly genes associated with neuronal activity and excitability suggesting a role for Gpr85 in modulating cerebellar GCs electrophysiological activity.

### Spontaneous activity of adult cerebellar Gpr85-KO GCs is increased *ex vivo*

Based on our transcriptomic analysis, we hypothesized that the electrophysiological properties of cerebellar GCs might be altered in Gpr85-KO models. To investigate this, we conducted electrophysiological recordings on *ex vivo* horizontal cerebellar slices from adult *gpr85*^GAL4/+^ (or *gpr85*^GAL4/GAL4^), *Tg(UAS:Lifeact-eGFP)* zebrafish (Fig. 6A). Recordings focused on *gpr85*-expressing GCs (eGFP^+^), to assess their spontaneous neuronal activity and passive membrane properties (Fig. 6A-C).

**Figure 6.**
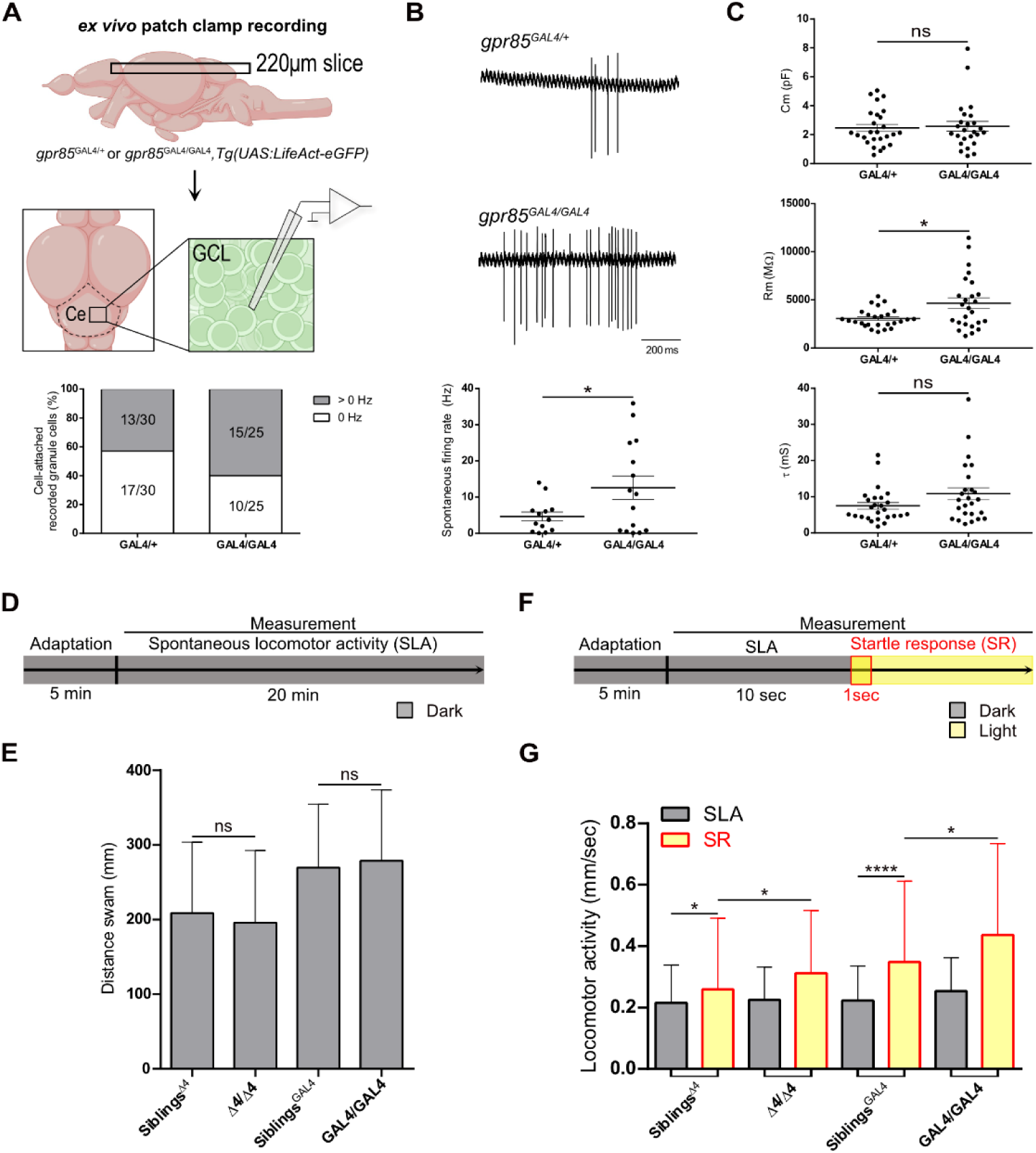
Gpr85-KO impacts adult cerebellar GCs electrophysiological properties and larval light-induced startle response. A Schematic representation of the experimental setup for recording the electrophysiological properties of adult gpr85-expressing cerebellar GCs *ex vivo*. Cerebellar horizontal sections (220 µm) were prepared, and GCs were identified based on GFP fluorescence. Recordings were made in cell-attached and whole-cell voltage clamp configurations. The chart illustrates the percentage of GCs in cell-attached configuration that exhibited or not spontaneous activity. B Quantification of GCs spontaneous activity in a cell-attached configuration (firing patterns are shown, as well as the firing rates measured over a 60-second recording (*gpr85*^GAL4/+^, n = 13 vs *gpr85*^GAL4/GAL4^, n = 15, *, p = 0.0347, unpaired t-test with Welch’s correction). C Quantification of GCs passive properties (membrane capacitance, Cm; membrane resistance, Rm; membrane time constant, τ) in a whole-cell voltage clamp configuration (*gpr85*^GAL4/+^, n = 26 vs *gpr85*^GAL4/GAL4^, n = 25; Cm, ns, p = 0.9132; τ, ns, p = 0.1724; Mann-Whitney U tests; Rm, *, p = 0.0113, unpaired t-test with Welch’s correction). D Schematic representation of the experimental setup for assessing spontaneous locomotor activity. The locomotor activity of larvae was recorded for 20 minutes in the dark after a 5-minute adaptation period. E Quantification of the distance swum by 6 dpf Gpr85-KO larvae compared to siblings over 20 minutes in the dark (*gpr85*^Δ4/Δ4^, n = 69 vs siblings, n = 210, p = 0.36; and *gpr85*^GAL4/GAL4^, n = 77, vs siblings, n = 190, p = 0.94, ns, not significant, Mann-Whitney U tests). F Schematic representation of the experimental setup for measuring light-induced startle responses. Larvae were recorded after a 5-minute habituation period. Spontaneous locomotor activity in the dark was recorded for 10 seconds as a baseline, followed by continuous light stimulation. The startle response was recorded within the first second following light onset. G The startle response was observed in controls as a significant increase in the distance swum following light onset (n = 210 siblings of *gpr85*^Δ4/Δ4^ mutants, dark vs startle response, *, p = 0.02; n = 247 siblings of *gpr85*^GAL4/GAL4^ mutants, dark vs startle response, ****, p<0.0001, Wilcoxon matched-paired signed-rank tests). The startle response was significantly more pronounced in *gpr85*-deficient animals compared to controls (n = 80 *gpr85*^Δ4/Δ4^ mutants vs siblings, *, p = 0.015; n = 83 *gpr85*^GAL4/GAL4^ mutants vs siblings, *, p = 0.011, Mann-Whitney U tests). Data information: Ce, cerebellum; GCL, granule cell layer; SLA, spontaneous locomotor activity; SR, startle response. Data are presented as mean ± SEM (B, C) or SD (E, G).

Cell-attached recordings revealed that in both genotypes, GCs displayed either no spontaneous activity (*gpr85*^GAL4/+^: 17/30, 57% vs *gpr85*^GAL4/GAL4^: 10/25, 40%) or some spontaneous activity (*gpr85*^GAL4/+^: 13/30, 43% vs *gpr85*^GAL4/GAL4^: 15/25, 60%) (Fig. 6A). Among spontaneously active neurons, the firing rate was significantly higher in Gpr85-KO GCs compared to controls (Fig. 6B, Hz, *gpr85*^GAL4/+^, 4.69 ± 1.24 vs *gpr85*^GAL4/GAL4^, 12.62 ± 3.24, p = 0.035, t-test). This result provides the first direct evidence that Gpr85 regulates the excitability of the adult cerebellar GCs.

We next investigated the passive membrane properties of neurons using a whole-cell patch voltage clamp configuration. While the capacitance (Cm) was not significantly different between our groups (Fig. 6C), we observed a significant increase in membrane resistance (Rm) in Gpr85-KO GCs (Cm: *gpr85*^GAL4/+^, 2.46 ± 0.25 pF vs *gpr85*^GAL4/GAL4^, 2.58 ± 0.34 pF, p = 0.913, M-W test; Rm: *gpr85*^GAL4/+^, 3079 ± 188 MΩ, n = 26 vs *gpr85*^GAL4/GAL4^, 4663 ± 554 MΩ, p = 0,011, t-test). This was not accompanied by significant changes in the membrane time constant (mS, *gpr85*^GAL4/+^, 7.47 ± 0.93 ms vs *gpr85*^GAL4/GAL4^, 10.81 ± 1.65 ms, p = 0.172, M-W test) (Fig. 6C).

Altogether, these results indicate that Gpr85 influences the intrinsic electrophysiological properties of adult cerebellar GCs. Specifically, its loss is associated with increased firing rates and higher membrane resistance, suggesting a role for Gpr85 in the regulation of neuronal excitability.

### Gpr85-KO larvae exhibit a stronger light-triggered startle response

We previously highlighted that Gpr85 is enriched at excitatory and inhibitory synapses in both the retina and the brain. We also reported that the receptor is expressed in key anatomical regions of the locomotor network and visual system, including the retina, optic tectum, cerebellum and spinal cord. Moreover, transcriptomic analyses of adult Gpr85-KO cerebellar GCs revealed gene expression changes related to neuronal activity and excitability. Finally, electrophysiological studies showed that Gpr85 deficiency alters the intrinsic neuronal properties of adult cerebellar GCs. Together, these findings led us to hypothesize that Gpr85 contributes to the regulation of neuronal activity in the locomotor and/or visual systems.

To test this, we conducted behavioral experiments on Gpr85-KO larvae using the Zebrabox system to evaluate locomotor responses to dark/light stimuli.

*Gpr85* heterozygous fish were incrossed, and their 6 dpf offspring were randomly and individually distributed into 48-well plates for analysis. We first recorded twenty minutes of activity in the dark to assess spontaneous locomotor activity under basal conditions (Fig. 6D). No significant differences in basal activity were observed between Gpr85-KO larvae and their siblings (*gpr85*^Δ4/Δ4^ vs siblings^Δ4^, p = 0.36 and *gpr85*^GAL4/GAL4^ vs siblings^GAL4^, p = 0.94, M-W test) (Fig. 6E).

We next assessed the light-triggered startle response of Gpr85-KO larvae by measuring the distance swam during the first second following light onset (Fig. 6F). In control larvae, this stimulus reliably evoked a startle response, as evidenced by a significantly longer distance swam after light stimulation compared to the dark condition (siblings^Δ4^: dark vs startle response, p = 0.02; siblings^GAL4^: dark vs startle response, p < 0.0001, Wilcoxon test) (Fig. 6G, Movie 1).

Both Gpr85-KO models displayed a significantly enhanced light-triggered startle response, reflected by a significantly increased distance swam following light induction as compared to their respective controls (*gpr85*^Δ4/Δ4^ mutants vs siblings^Δ4^, p = 0.015; *gpr85*^GAL4/GAL4^ mutants vs siblings^GAL4^, p = 0.011, M-W test) (Fig. 6G). These results suggest that Gpr85 plays a role in modulating neuronal responses to light stimuli and contributes to the regulation of visual and locomotor system activity.

## Discussion

Although GPR85 was identified twenty-five years ago as one of the most well-conserved receptors in vertebrates, its biological functions remain elusive. Here, we significantly expanded the understanding of *gpr85* expression and function. We provide the first evidence that Gpr85 is enriched at the plasma membrane of neuronal chemical synapses *in vivo* and demonstrate that constitutive Gpr85 deficiency alters the activity of adult zebrafish cerebellar GCs and light-induced motor behavior in larvae.

Using a new reporter line and single-cell transcriptomics, we mapped *gpr85* expression across development and adulthood, confirming its presence in the brain, retina, spinal cord, and intestine. Adult expression was detected in brain regions conserved across species—olfactory bulb, pallium, habenula, hypothalamus, cerebellum, and optic tectum—as well as in the testis and intestinal neurons, aligning with human and mouse data (Matsumoto et al., 2000) (Hellebrand et al., 2001). A novel finding was its expression in developing neurons of the larval intestine.

At the subcellular level, we demonstrate that Gpr85 is enriched at excitatory and inhibitory synapses, and at ribbon synapses in the retina, consistent with prior in vitro observations of PSD-95 binding (Fujita-Jimbo et al., 2015) and synaptosomal enrichment (Yoshimura et al., 2004). In the developing and adult retinal IPL, Gpr85 was also enriched at the plasma membrane of ribbon synapses, at both pre- and postsynaptic sites. This observation is consistent with our previous observation that GPR85 is targeted to the plasma membrane *in vitro* (Kaafarani et al., 2023). Interestingly, when detected on both sides of ribbon terminals, the receptor exhibited a reciprocal, clustered distribution, suggesting non-uniform localization within synapses.

Unlike the sole documented GPR85-KO mouse model (Matsumoto et al., 2008; Chen et al., 2012), our zebrafish Gpr85-KO models exhibit no brain mass defects. Developmental synaptogenesis in *gpr85*-expressing neurons appeared unaffected. However, single-cell RNA-seq revealed that Gpr85-KO cerebellar GCs showed strong transcriptional changes associated with excitability—including upregulation of immediate early genes (IEGs), ion channels, and synaptic vesicle components. This supports a model where Gpr85 modulates neuronal activity rather than development.

Notably, *oaz2b* and *oaz1a*, regulators of polyamine synthesis, were among the most upregulated genes. Given the known links between polyamines and neuronal excitability (Pegg, 2009), and their dysregulation in psychiatric conditions (Guipponi et al., 2009), this raises the possibility that Gpr85 influences GC physiology through the polyamine pathway.

Electrophysiological recordings of adult GCs—performed for the first time in zebrafish— confirmed that Gpr85-KO cells are more spontaneously active and exhibit increased membrane resistance. Notably, while mammalian GCs are typically characterized by either no spontaneous activity *ex vivo* (D’Angelo, 2013) or a low firing rate *in vivo* (∼0.5Hz) (Chadderton et al., 2004), we recorded GCs with no (57%) or with a spontaneous activity in our controls (43%, 4.69 ± 1.24 Hz). Zebrafish cerebellar neuronal populations and the organization of the granule cell layer glomeruli are well conserved compared to mammals (Bae et al., 2009; Pose-Méndez et al., 2023). Glomerular synapses are both excitatory and inhibitory, mainly involving phasic glutamatergic and GABAergic transmission (D’Angelo, 2013). Given the thickness of our horizontal cerebellar slices (220 µm, approximately half the thickness of the adult zebrafish granule cell layer) and the absence of synaptic transmission blockage, the recorded activity is potentially influenced by residual inhibitory and/or excitatory synaptic transmission.

Consistent with the passive properties of rat cerebellar GCs (Chadderton et al., 2004), voltage-clamp assessments revealed that adult zebrafish cerebellar GCs exhibit a low membrane capacitance (2.46 ± 0.25 pF) and an average membrane time constant of 7.5 ± 0.93 ms. Further analysis revealed an increase in the membrane resistance (Rm) of Gpr85-KO GCs, while the membrane time constant and capacitance remained unaffected. This suggests fewer active and/or less abundant ion exchangers at the plasma membrane at −60mV. In our transcriptomic study, we observed upregulation of *ATPases Na^+^/K^+^* pump subunits, including *ATPase Na^+^/K^+^ transporting subunit alpha 3b/1a.1* and *beta1b/2a*. These pumps play a major role in establishing the resting membrane potential (Pivovarov et al., 2018), suggesting that this upregulation likely represents a compensatory adaptation rather than a direct cause of the elevated Rm. While the mechanism by which Gpr85 affects Rm and the firing rate requires further investigation, our transcriptomic and electrophysiological findings suggest that Gpr85 modulates intrinsic properties and excitability of adult cerebellar GCs. Despite the recording of action potentials during whole-cell current-clamp protocols, the access resistance exceeded the standard limits, precluding reliable assessment of adult zebrafish GCs excitability *ex vivo*.

Functionally, Gpr85-KO larvae displayed enhanced light-triggered startle responses, consistent across two independent KO models. This phenotype, though subtle, supports a role for Gpr85 in regulating sensory responsiveness, potentially via visual or cerebellar circuits. Our findings align with behavioral alterations reported in GPR85 mutant mice (Matsumoto et al., 2008). In line with this result, GPR85 knock-down and overexpression approaches have suggested that the receptor contributes to bone cancer pain in the rat spinal cord (Ni et al., 2023).

Although the precise mechanisms linking Gpr85 deficiency to the functional alteration of cerebellar GCs and behavioral changes of fish remain to be delineated, a number of potential links can be underlined with the sets of genes whose expression is modified in Gpr85-KO GCs. IEGs are well established markers of neuronal activity, including in GCs (Monti et al., 2002; Contestabile et al., 2005), and their upregulation likely reflects their increased excitability. The upregulation of the *oaz* genes in Gpr85-KO GCs may lead to polyamine depletion. This latter is known to enhance NMDA receptor activity and alter AMPA and potassium channel functions, thereby increasing GC excitability (Makletsova et al., 2022). Finally, our findings may have broader implications given the association of GPR85 with autism spectrum disorders (ASD) (Fujita-Jimbo et al., 2015; Wu et al., 2022). ASD encompasses a range of neurodevelopmental diseases characterized by deficits in communication and social interactions, as well as repetitive behaviors (Lord et al., 2018). There is mounting evidence pointing to a role of the cerebellum in cognition and emotion (Ma et al., 2023). Cerebellar GCs play key roles in motor coordination, cognitive processing and sensory integration (Rudolph et al., 2023), and their functional alteration, as observed in our Gpr85-KO fish, might lead to an imbalance of cerebello-cortical circuits leading to cognitive and social deficits observed in ASD.

In conclusion, our study shows that Gpr85 is localized at chemical synapses *in vivo*, and influences adult cerebellar granule cells electrophysiological activity as well as light-induced motor behavior in zebrafish larvae. These findings establish a foundation for exploring Gpr85’s role in vertebrate brain function and its potential contribution to neurodevelopmental disorders.

## Material and methods

### Zebrafish husbandry

Zebrafish (AB* strain) were raised and maintained under standard laboratory conditions, in accordance with the Federation of European Laboratory Animal Science Associations (FELASA) guidelines (Aleström et al., 2020). All experimental procedures were approved by the Ethical Committee for Animal Welfare (CEBEA) of the Faculty of Medicine, Université Libre de Bruxelles. The zebrafish mutants and transgenic lines used in this study were *Tg(UAS:GFP-CAAX)*^m1230^ (Fernandes et al., 2012), *Tg(UAS:lifeact-eGFP)*^mu271^ (Helker et al., 2013) and *Tg(Xltubb:DsRed)*^zf148^ (Peri & Nüsslein-Volhard, 2008), here referred to as *Tg(NBT:DsRed)*. The term “adult” fish refers to animals aged between 6 and 8 months of age. For consistency and clarity throughout the text, transgenic animals are referenced without their allele designations.

### Generation of *gpr85*^Δ4^ knock-out and *gpr85*^GAL4^ knock-in mutants

Two sgRNAs were designed using Sequence Scan for CRISPR and CRISPR Scan software (http://crispr.dfci.harvard.edu/SSC/ and http://www.cirsprscan.org/) (Xu et al., 2015). The first *gpr85*-targeting sgRNA1, used to generate the *gpr85*^Δ4^ line, was synthetized *in vitro* as described previously (Talbot & Amacher, 2014). *Gpr85*^GAL4^ mutants were generated using the Geneweld strategy, a variant of CRISPR/Cas9 technology for targeted integration (Wierson et al., 2020; Jordan M et al., 2021). The GAL4 coding sequence was inserted in frame with the endogenous *gpr85* start codon. We co-injected 0.5 nL of a solution containing Cas9 (100 ng/µL), the *gpr85*-targeting sgRNA2 (70 ng/µL), UgRNA (40 ng/ µL) and the GAL4 donor vector (30 ng/µL) into *Tg(UAS:lifeact-eGFP)* one-cell stage embryos. According to the Geneweld approach, the GAL4 cargo was flanked by UgRNA-targeted sequences and 48 bp homology arms. F0 founders were screened at 3 dpf based on eGFP fluorescence. For both lines, individuals from at least the F3 generation were used in experiments. Mutations were validated by PCR and Sanger sequencing (Eurofins). Specific guide RNAs, primers and homology arms used are listed in the Reagents and tools table (Table 1).

### Generation of transgenic lines

The following transgenes were randomly integrated into the genome of the *gpr85*^GAL4^ line: *Tg(zcUAS:FinGR(PSD95)-GFP-ZFC(CCR5TC)-KRAB(A))*, *Tg(ziUAS:FinGR(GPHN)-mCherry-ZFI(IL2RGTC)-KRAB(A))*, *Tg(UAS:zGpr85-eGFP), Tg(UAS:zGpr85-mCherry).* To achieve this, we co-injected 0.5 nL of a solution containing 120 ng/µL of *tol2* transposase mRNA and 40 ng/µL of the respective *tol2*-based vector into one-cell stage embryos. The *tol2*-UAS:zGpr85-eGFP vector was generated by digesting and ligating a *tol2*-UAS-MCS-EGFP plasmid (Genecust, Puc57 customed) with the amplified zebrafish *gpr85* coding sequence using *Xho*I and *Bbv*CI restriction sites. The eGFP cassette of this newly generated vector was then replaced by an mCherry cassette using *Bbv*CI and *Sal*I restriction sites in order to generate the *tol2*-UAS:zGpr85-mCherry construct. The pTol2-zcUAS:PSD95.FinGR-GFP-ZFC(CCR5TC)-KRAB(A) and pTol2-ziUAS:GPHN.FinGR-mCherry-ZFI(IL2RGTC)-KRAB(A) plasmids were obtained from Addgene (plasmids #72638 and #72639, respectively). F0-injected embryos were screened and selected based on their GFP or mCherry fluorescence. Each F1 progeny derived from an F0 founder was screened for fluorescence and incrossed to establish stable F2 lines. For clarity, *Tg(zcUAS:FinGR(PSD95)-GFP-ZFC(CCR5TC)-KRAB(A))* and *Tg(FinGR(GPHN)-mCherry-ZFI(IL2RGTC)-KRAB(A))* lines are further referred to in the text as *Tg(UAS:FinGR(PSD)-GFP)* and *Tg(UAS:FinGR(GPHN)-mCherry)*, respectively.

### WISH

Sense and antisense riboprobes for *gpr85* (ENSDARG00000068701) were synthesized *in vitro* using cDNA from 3 dpf embryos. PCR primers used are listed in in the Reagents and tools table (Table 1). Forward or reverse primers contain the T3 polymerase promoter sequence. WISH was performed using a standard protocol (Thisse & Thisse, 2008) with a minimum of 20 individuals per condition. To ensure specificity of the staining, the sense and antisense riboprobes were always hybridized in parallel. Probe hybridization was carried out at 65°C and BM-purple (Roche) was used as the substrate for signal revelation.

### Gene expression analysis by qPCR

For each biological replicate, 30 larvae were digested in TRIzol reagent (Qiagen). RNA was isolated by chloroform extraction and purified with the RNAeasy Mini Kit (Qiagen). Genomic DNA contamination was removed using the TURBO DNA-*free* kit (Thermo Fisher Scientific). The cDNAs were prepared from 2 µg of purified RNA using Super Script II reverse transcriptase with Oligo(dT) primers (Thermo Fisher Scientific). qPCRs were performed using the CFX96 Real-Time System (Bio-rad, CA) according to the manufacturer’s instructions. Biological replicates were tested in duplicate, and relative transcript levels were assessed using the ΔC_T_ method. Gene expression levels were normalized to two reference genes (*rpl13* and *lms12b*). Primers used are listed in the Reagents and tools table (Table 1).

### Fluorescent immunostaining

Embryos and larvae were fixed overnight in 4% PFA and processed for either whole-mount or cryosection antibody staining. Adult organs were dissected and fixed in 4% PFA at 4°c for 4 hours, then incubated for over 2 days at 4°c in 30% sucrose. The samples were then embedded in OCT, snap-frozen and cryosectioned (14 µm, Leica CM3050 S). Staining procedures were performed according to previously described protocols (whole-mount: Ferrero et al., 2018; cryosections: Ferrero et al., 2020). A minimum of 3 individuals were imaged for each condition. The primary and secondary antibodies used are listed in the Reagents and tools table (Table 1).

### Zebrafish morphometric measurements

*Gpr85^GAL4^* and *Gpr85^Δ4^* heterozygous mutants were respectively incrossed and their corresponding adult offspring were analyzed. The body length was measured using a ruler. Brains were dissected, carefully dried using absorbent paper, and weighted using a precision balance (Sartorius Entris II, BCE224I-1S). Brains with dissection defects were discarded. All measurements were performed before genotyping.

### Tissue clearing

For adult brain tissue clearing, a modified EZ Clearing method (Hsu et al., 2022) was used. Zebrafish heads were dissected and enucleated, followed by overnight fixation in 4% PFA (pH 8.5) at 4°C. All subsequent steps were performed under agitation at 4°C. Whole fixed brains were dissected and incubated overnight in 50% (v/v) THF (Sigma-Aldrich, 186562) in sterile Milli-Q water (pH adjusted to 8.5 with triethylamine, Sigma-Aldrich, T0886). Samples were then washed four times (1h each) with sterile Milli-Q water. Finally, samples were submerged in refractive index (RI) matching EZ View solution (80% Nycodenz, Accurate Chemical & Scientific AN1002423, 7 M Urea, 0.05% sodium azide in 0.02 M sodium phosphate buffer; prepared according to Hsu et al., 2022) and stored at 4°C for at least 24h before imaging.

### Electron microscopy

#### Structural analysis

Wild-type (WT) larvae were fixed for 4 days at 4°C in 2.5% glutaraldehyde (Sigma-Aldrich). Unless specified, all subsequent steps were performed at room temperature. The samples were rinsed in 0.1 M cacodylate buffer, pH 7.4, then stained for 1 h in 1% osmium tetroxide-1.5% ferrocyanide (0.1 M cacodylate), 1 h in 1% osmium tetroxide (0.1 M cacodylate), followed by 1.5 h in 1% uranyl acetate (ultrapure water). The larvae were next embedded in 1% agarose and underwent dehydration in graded ethanol solutions (50%, 70%, 95% and 100%). Ethanol was then replaced with 100% propylene oxide (PO, two successive 8-min baths), then with a 50/50 mixture of PO/epoxy resin (Agar scientific, Agar 100) for 1 h, and with 100% epoxy resin for 3 h. The resin was finally changed and polymerized for 2 days at 60°C. The resulting blocks were sectioned using a UC7 ultramicrotome (LEICA EM) and collected on carbon-formvar 100 mesh copper grids (Electron Microscopy Sciences).

#### Immunogold staining

*Gpr85*^GAL4/+^; *Tg(UAS:zgpr85-mCherry)* larvae were fixed for 2 weeks at 4°C in 4% PFA-0.5% glutaraldehyde, then embedded in 12% gelatin blocks (in ultrapure water). Unless specified, the following steps were performed at room temperature. Microdissection was performed to retain only the head of the zebrafish. The obtained blocks were incubated for 2 days in 2.3 M sucrose on a rotating wheel at 4°C, then stored at 4°C before processing. Next, the blocks were mounted on supports and sectioned using a UC6 ultramicrotome equipped with an FC6 cryo-chamber (LEICA EM). Sections were made at −120°C and placed on carbon-formvar 100 mesh nickel grids (Electron Microscopy Sciences). The grids were processed for immunogold labeling as follows. Samples were blocked for 30 minutes in 5% goat serum and incubated O/N at 4°C with the following primary antibodies: rabbit anti-DsRed polyclonal (632496, Takara, 1:50) or rabbit polyclonal anti-Ribeye-A (s4561-2, Zenisek’s lab, 1:50) in 3% goat serum. The grids were washed in TBS buffer, then incubated with a secondary anti-rabbit antibody (1:100, Sigma-Aldrich) for 1h at 37°C in TBS. The grids were washed in TBS buffer, followed by washes in ultrapure water. Finally, the grids were stained for 10 minutes on ice with methylcellulose-4% uranyl acetate (9:1). The grids were observed using a Tecnai 10 100kV transmission electron microscope (FEI, Thermo Fisher Sci). Images were captured with a MegaView 14 bits camera (Olympus) and processed using the iTEM software (Olympus). Three individuals were used for imaging.

#### Light microscopy imaging and image analysis

WISH and live embryos/larvae were imaged using a Leica M165FC fluorescent stereomicroscope equipped with a Leica DFC7000T digital camera. Images were acquired using the LAS software (Leica, V4.6.2). Immunostained whole-mount larvae and cryosections were imaged using a Zeiss LSM 780 inverted confocal microscope, with Plan Apochromat 20x/0.8 M27, LD C Apochromat 40x/1.1 W korr M27 and Plan achromat 63x/1.46 oil Korr M27 objectives. Quantification of GFP^+^ neuronal soma within the optic tectum stratum paraventriculare was performed using ImageJ software. The area of individual hemisphere used for the quantification was drawn manually and the entire sections were analyzed (z-stacks, 3 µm steps). Quantification of FinGR(PSD)-GFP^+^ puncta density in the optic tectal neuropils was performed using Imaris software. Puncta detection was based on an estimated diameter of 0.3 µm with background subtraction applied. Two identical rectangular soma-free areas (1616 µm^2^ each) per optic section were defined for quantification. Three optic sections per hemisphere per larva were measured and averaged. Cleared brain samples were sagitally cut between the hemispheres and immersed cut-face down in a RI matching solution. Images were acquired with a Nikon AX/R confocal microscope system using a 10X/0.45 Plan Apo LambdaD objective lens in resonant scanning mode. Acquisition settings were adjusted to 1.3-1.8X zoom with 2048×2048 pixel resolution and a tile scan setting with 5% overlap. Online stitching mode was enabled in for real-time stitching during acquisition. Images were post-processed with 3D Deconvolution, and with Denoise.ai, in NIS-Elements AR software to obtain sharper Z-resolution. The whole brain hemispheres of a male and a female were imaged.

### Comparative scRNAseq and data analysis

#### scRNAseq library preparation and sequencing

Female adult brains from siblings (one control and one *gpr85*-KO) were dissected and digested with 14 U papain (Sigma-Aldrich, p4762) at 33°C for 10 to 15 minutes in 0.9X Dulbeccoʹs Phosphate Buffered Saline (DPBS). Mechanical dissociation was performed during this process using a syringe with a 26 G needle (B.Braun, Omnifi× 100 Duo). Cell suspensions were washed and centrifuged (300 g) in 2% fetal bovine serum diluted in 0.9X DPBS (FACS buffer). Then, cells were resuspended in FACS buffer and filtered using a sterile 40 µm nylon mesh (VWR). Cell sorting of eGFP^+^ cells was performed on a FACS ARIA (Becton Dickinson, San Jose, CA) using Sytox Red (5 nM, Thermofisher scientific S34859) in order to remove dead cells. 50,000 cells were sorted per condition with an expected final density of 294 cells/µL. The library was prepared following 10x Genomics Chromium Single cell 3’ kit (v3) guidelines and the libraries were sequenced using an Illumina NovaSeq 6000.

#### scRNAseq data processing

Raw sequencing data were processed using Cell Ranger (v7.1.0) with a custom-built reference based on the zebrafish reference genome GRCz11 and gene annotation Ensembl 92 in which the GFP sequence was added. Data were analyzed through the Seurat Package in R (Stuart et al., 2019), keeping only cells having between 500 to 10,000 counts with less than 7.5% of genes coming from the mitochondrial genome. SCTransform was used as the scaling method (Hafemeister & Satija, 2019). The UMAPs were generated using 10 dimensions with a clustering resolution of 0.5.

### Electrophysiology

Adult brains from *gpr85*^GAL4/+^ (or *gpr85*^GAL4/GAL4^), *Tg(UAS:Lifeact-eGFP)* individuals were dissected (n=3 per condition), and *ex vivo* horizontal cerebellar slices (220 µm thick) were produced using a Vibratome R VT 1000 S (Leica) in an ice-cold solution (2.5 mM KCl, 1.25 mM NaH_2_PO_4_, 25 mM NaHCO_3_, 7 mM MgCl_2_, 0.5 mM CaCl_2_, 14 mM glucose, and 139 mM choline chloride) gassed with a carbogen solution (95% O_2_ and 5% CO_2_). Then, slices were transferred to the recovery chamber at 26°C and incubated in artificial cerebrospinal fluid (aCSF) containing: 127 mM NaCl, 2.5 mM KCl, 1.25 mM NaH_2_PO_4_, 1 mM MgCl_2_, 26 mM NaHCO_3_, 10 mM D-glucose, 2 mM CaCl_2_, bubbled with a mixture of 95% O_2_ and 5% CO_2_ at a pH of 7.3 (300-316 mOsm). After a recovery period of 45 min, individual slices were transferred to the recording chamber continuously superfused with aCSF at a rate of 1-1.5 ml/min at room temperature (24-26°C). Cerebellar GCs were identified with a 63x water immersion objective from a Zeiss Axioskop microscope (Axioskop 2FS Plus; 140 Zeiss) equipped with an infrared CCD camera (XST70CE, Hamamatsu Photonics KK). GCs were selected based on their eGFP expression using the filter set 38 HE (BP: 470/40 wavelength, Carl Zeiss Instruments) and the OptoLED R electroluminescent diode (Cairn Research Lda). Borosilicate-glass patch electrodes (GmbH) (resistance between 5 and 7 MΩ) filled with 125 mM KMeSO_3_, 12 mM KCl, 0.022 mM CaCl_2_, 4 mM MgCl_2_, 10 mM HEPES, 0.1 mM EGTA, 5 mM Na_2_-phosphocreatine, 4 mM Mg_2_-ATP and 0.5 mM Na_2_-GTP, were used for the recordings. Cerebellar GCs were first recorded in cell-attached and then in whole-cell configurations to examine their intrinsic properties. Recordings were performed using an EPC-10 patch clamp amplifier (HEKA) and PatchMaster acquisition software (HEKA). Signals were sampled at 20 kHz with a gain of 2 mV/pA and lowpass filtered at 2.9 kHz.

Extracellular spontaneous action potentials were acquired in cell-attached configuration (Seal > 1 GΩ, 60 to 120 s recordings). Standard off-line detection of action potential was performed with Axograph X software (Axon Instruments Inc.). For this analysis, we generated an action potential template to scan the recording trace. All matching events were stored, and false positive events were detected and discarded based on their amplitude. Spontaneous firing frequencies were assessed with 60 second recordings. Cells were then recorded in whole-cell configuration in voltage-clamp mode at holding potential of −60 mV. Passive membrane properties and access resistance were extracted from current traces, averaged on ten sweeps recorded in response to a hyperpolarizing voltage pulse (200 ms) of 10 mV from holding potential. The membrane resistance was computed using the difference between the baseline current and the current at 20 ms during the voltage step. The integrated area under the transient was used to determine membrane capacitance. Membrane time constant was calculated as the product of the Rm multiplied by the Cm. If access resistance changed more than 25% between the beginning and the end of the recording, the neuron was excluded from the analysis. All analysis were performed using IgorPro 6.3 software (WaveMetrics) using Patcher’s Power Tools, NeuroMatic plugins and Microsoft Excel software were used.

### Behavioral experiments

Larvae were randomly assigned and individually immerged in 1 mL of E3 standard medium (5 mM NaCl, 0.17 mM KCl, 0.33 mM CaCl_2_, 0.33 mM MgSO_4_, dH_2_O) (Fetcho, 2003) in 48 well-plates (VWR) and placed in a standard Zebrabox (Viewpoint) for recording. Larvae presenting developmental defects such as trunk curvature and/or pericardial edema were excluded from the experiments. The temperature of the chamber was maintained at 28°c and all recordings were performed after a 5-min habituation step in the dark. Experimental set up and data were generated using Viewpoint’s Zebralab software. The spontaneous locomotor activity in the dark was assessed over a 20-min integration period while the light-triggered startle response was induced using 100% light intensity (8000 lx) and measured over a 1-second integration period.

### Statistical Analysis

Statistical analyses were conducted using Graphpad Prism8. Normality was assessed using the D’Agostino-Pearson omnibus normality test and Student’s t-test or Mann-Whitney U test was applied accordingly. Wilcoxon matched-pairs rank test was used to compare basal locomotor activity in the dark versus light-induced startle response for a given condition. The results are considered significant when p < 0.05 and are displayed with the mean and standard error of the mean (SEM) or the standard deviation (SD). Data display and sample sizes used for the statistical analysis are specified in the respective figure legends.

## Supporting information

Data set1

Reagents and tools table1

## DATA availability

The datasets produced in this study are available at GSE291414.

## Acknowledgements

We are grateful to Sumeet Pal Singh, Elif Sema Eski, and Gilles Dinsart for their precious advices and their assistance with single-cell data processing. We thank Jean-Marie Vanderwinden and Michiel Martens from the ULB Limif Platform, as well as Christine Dubois from the FACS facility. We thank Courtney Elaine Frederick and David Zenisek for generously donating an aliquot of their αRibeye antibody. We thank Jeffrey Farrel and Abhinav Sur for their permission to use “Daniocell’s” website images. We also thank Eva Laurrel and the other members of the zebrafish “mapzebrain” atlas project. Finally, we thank Mustapha Chaouni and Kamel Gharbi for their daily maintenance of the fishroom facility. This project was funded by the “Fonds David et Alice Van Buuren” and the Fonds National de la Recherche Scientifique (FNRS: CDR J021.517F and FRIA fellowship).

## Conflict of interest statement

All authors declare no conflict of interest.

## Author contributions

Romain Darche-Gabinaud: Conceptualization; Data curation; Formal analysis; Funding acquisition; Investigation; Visualization; Methodology; Writing—original draft; Writing— review and editing.

Abeer Kaafarani: Conceptualization; Molecular biology; Review and editing. Marine Chazalon: Resources; Data curation; Formal analysis; Investigation.

Valérie Suain: Resources; Data curation; Formal analysis; Investigation; Visualization; Methodology.

Erika Hendrick: Resources; Data curation; Formal analysis; Investigation; Visualization; Methodology.

Louise Conrard: Resources; Data curation; Formal analysis; Investigation; Visualization; Methodology.

Anne Lefort: Resources; Formal analysis; Methodology.

Frédérick Libert: Resources; Software; Formal analysis; Methodology.

Mehmet Can Demirler: Resources; Visualization; Methodology.

Serge Schiffmann: Conceptualization; Resources; Data curation; Supervision; Writing— Review and editing.

David Perez-Morga: Conceptualization; Resources; Data curation; Supervision.

Valérie Wittamer: Conceptualization; Resources; Data curation; Software; Formal analysis; Supervision; Funding acquisition; Validation; Investigation; Visualization; Methodology; Project administration.

Marc Parmentier: Conceptualization; Resources; Data curation; Software; Formal analysis; Supervision; Funding acquisition; Validation; Investigation; Visualization; Methodology; Writing—original draft; Project administration; Writing—review and editing.

Isabelle Pirson: Conceptualization; Resources; Data curation; Software; Formal analysis; Supervision; Funding acquisition; Validation; Investigation; Visualization; Methodology; Writing—original draft; Project administration; Writing—review and editing.

---

**Figure 1 – figure supplement 1.**
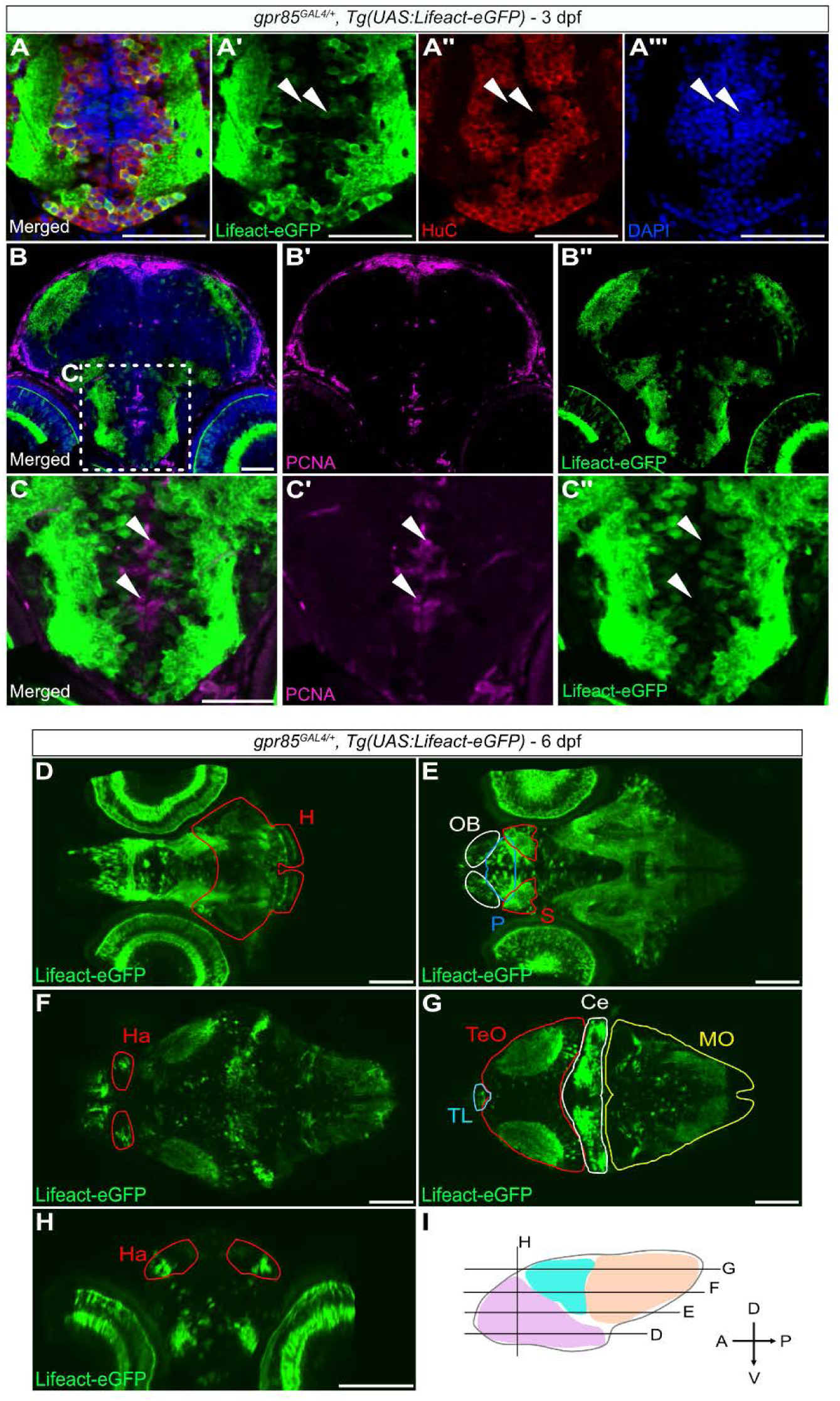
The *Gpr85* reporter line shows non-proliferating cells expressing *gpr85* in the developing brain mesenchyme, and *Gpr85* is broadly expressed in the post-embryonic developing brain (mapzebrain atlas). A Confocal imaging of the midbrain ventral region of 3 dpf *gpr85*^GAL4/+^*, Tg(UAS:Lifeact-eGFP)* zebrafish larvae stained with anti-GFP (green), anti-HuC (red), and DAPI (blue). Some Lifeact-eGFP^+^ cells can be observed in the HuC^-^ neural progenitor/precursor area (white arrowheads). Scale bar: 50 µm. B Maximum projection confocal images of coronal sections of the brain of 3 dpf *gpr85*^GAL4/+^*, Tg(UAS:Lifeact-eGFP)* zebrafish larvae stained with anti-GFP (green), anti-PCNA (magenta), and DAPI (blue). The box indicates the region enlarged in panel C. Scale bar: 50 µm. C Higher magnification of the boxed region in panel B, focusing on the ventral telencephalon. None of the PCNA^+^ cells in the progenitor/precursor zone are Lifeact-eGFP^+^ (white arrowheads). Scale bar: 50 µm. D-G Confocal images of horizontal sections from a 6 dpf *gpr85*^GAL4/+^*, Tg(UAS:Lifeact-eGFP)* zebrafish larva showing Lifeact-eGFP staining in the highlighted anatomical regions. Full dataset is available online on https://mapzebrain.org/atlas/2d (marker: *gpr85*; Shainer et al., 2023). Scale bars: 100 µm. H Confocal image of a coronal section of a 6 dpf *gpr85*^GAL4/+^*, Tg(UAS:Lifeact-eGFP*) zebrafish larva showing Lifeact-eGFP^+^ cells in the habenula. Scale bar: 100 µm. I Schematic representation of the shown sections through a 6 dpf zebrafish brain, with the forebrain in pink, the midbrain in turquoise, and the hindbrain in peach. Data information: Ce, cerebellum; dpf, days post fertilization; Ha, habenula; H, hypothalamus; MO, medulla oblongata; OB, olfactory bulb; P, pallium; S, subpallium; TeO, optic tectum; TL, torus longitudinalis.

**Figure 2-figure supplement 1.**
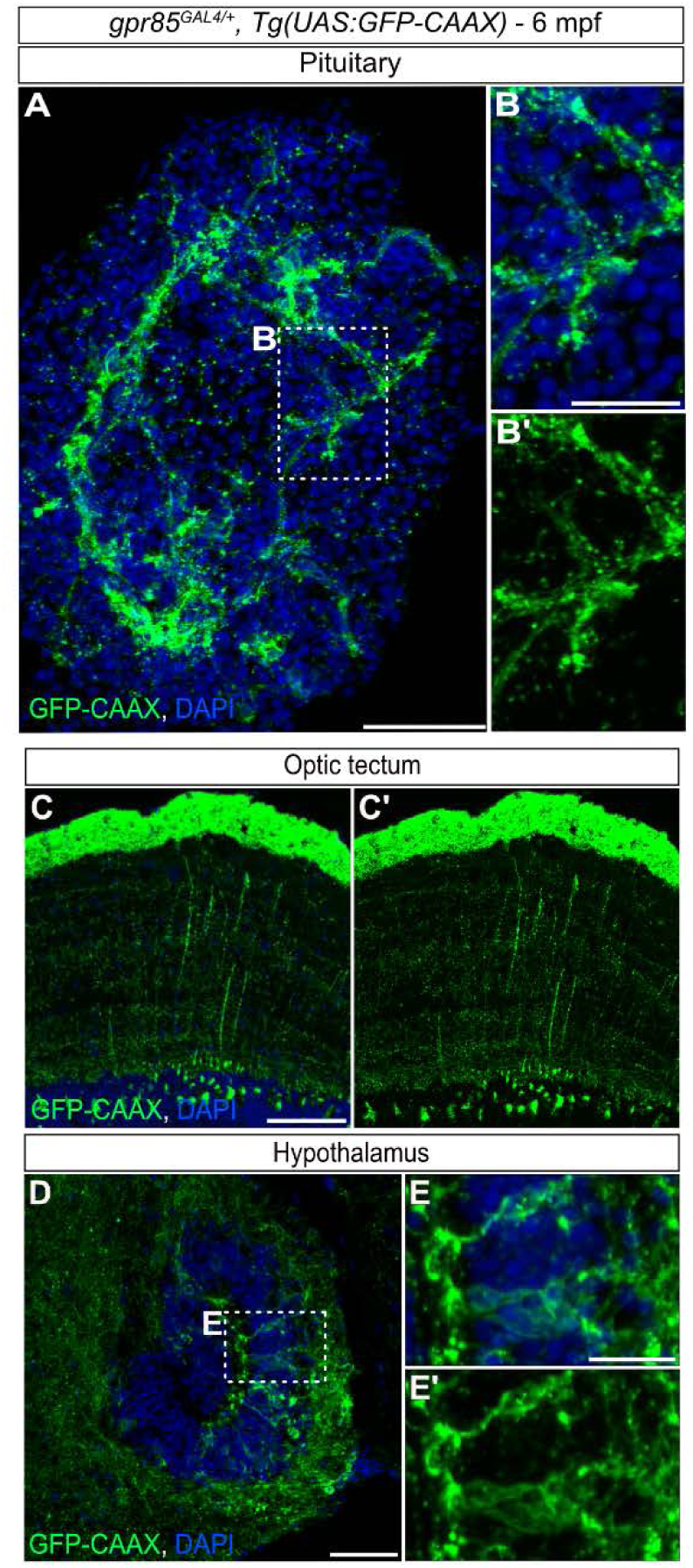
Cartography of *gpr85* expression in the pituitary, hypothalamus and optic tectum of adult zebrafish. A-E’ Maximum projection confocal images of brain regions from a *gpr85*_GAL4/+_*, Tg(UAS:GFP-CAAX)* adult zebrafish stained with anti-GFP (green) and DAPI (blue). (A) Coronal section of the pituitary gland. The box indicates the region enlarged in panel B. (B) Enlarged view of the region boxed in A. (C-E’) Sagittal sections of the optic tectum (C) and the caudal hypothalamus (D). The box in D indicates the regions enlarged in panels E. (E) Enlarged view of the region boxed in D. Scale bars: (A, C, D) 50 µm, (B, E) 25 µm.

**Figure 5-figure supplement 1.**
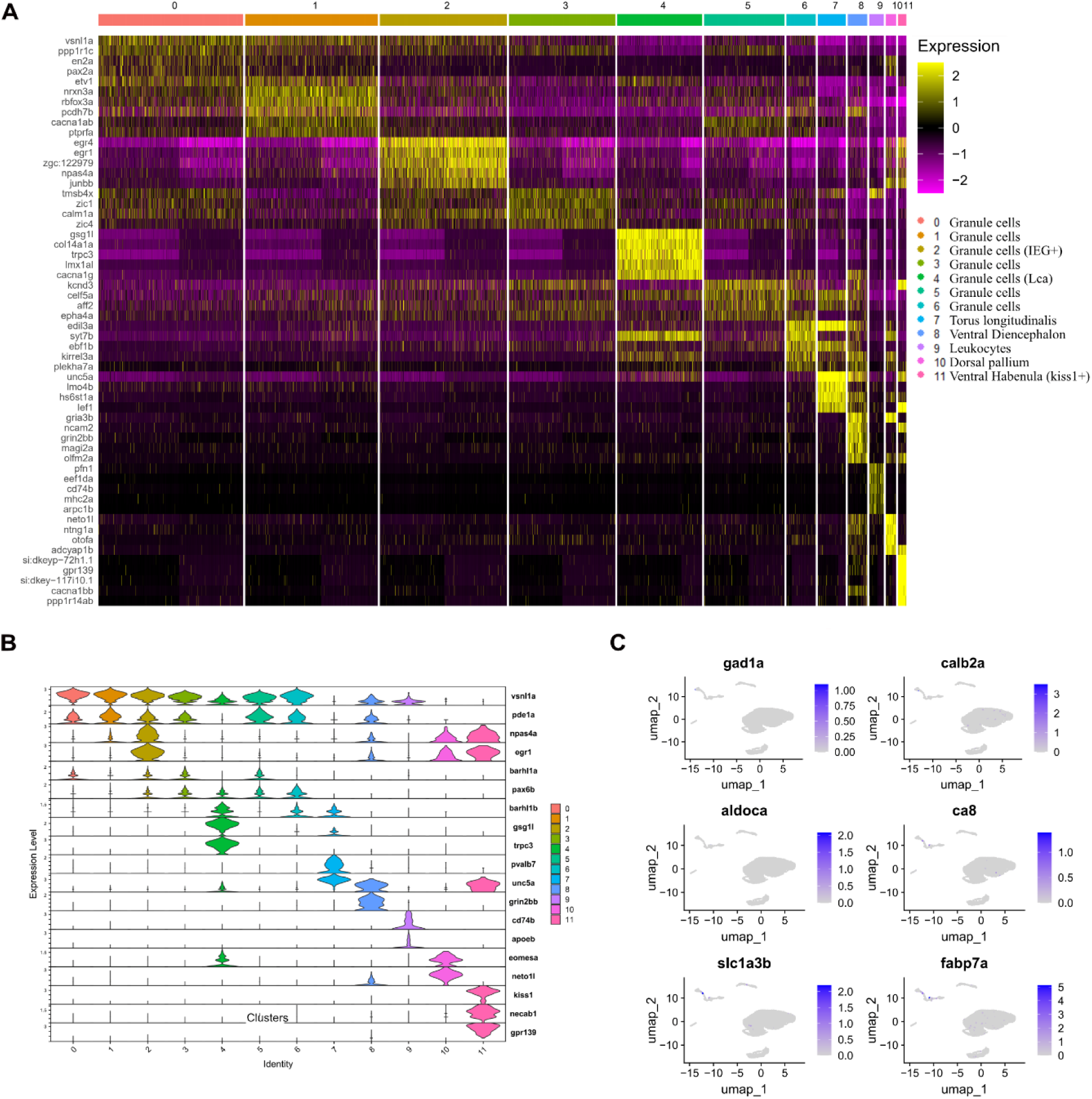
scRNAseq supplementary: clustering markers. A Heatmap of the top 5 markers for each individual cluster. All cells are represented. B Violin plots of relevant markers used for the identification of the different clusters. C UMAPs of the Ctl and KO cells merged, displaying the expression of key cerebellar markers used to discriminate eurydendroid, Purkinje, stellate, and Golgi cells.

## Notes

### Competing Interest Statement

The authors have declared no competing interest.

### Summary of Updates

This version of the manuscript has been revised to update the abstract and the introduction in order to clarify the message and the discussion to shorten it.

