## Supplementary Figures and Table for "Synaptic Gpr85 influences cerebellar granule cells electrical properties and light-induced behavior"

**Table of content**

Appendix Figures: page 2-6

Appendix Table: page 7

Appendix Figures Legends: page 8-11

Appendix Tables Legends: page 12

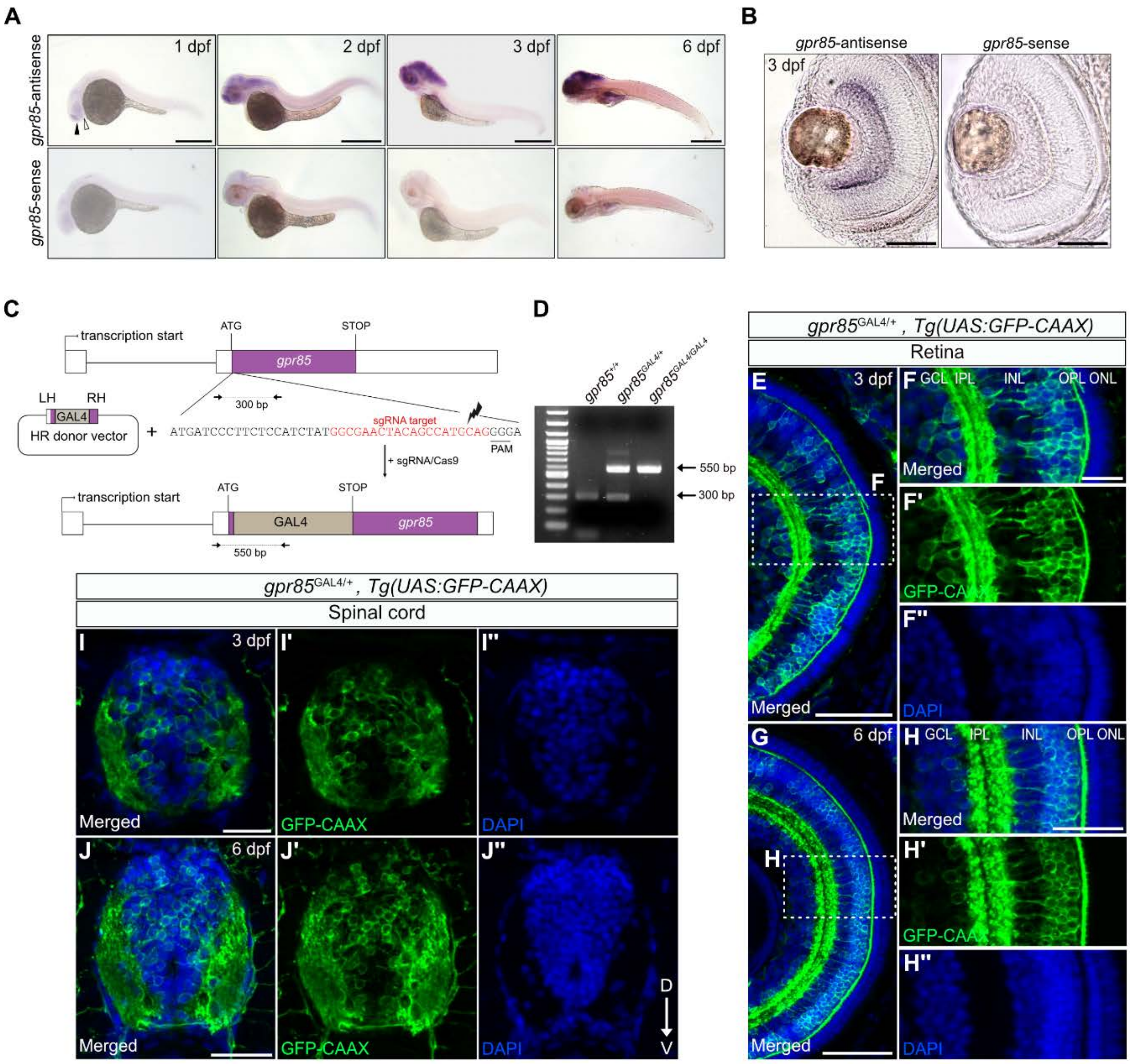


### Appendix Figure S1

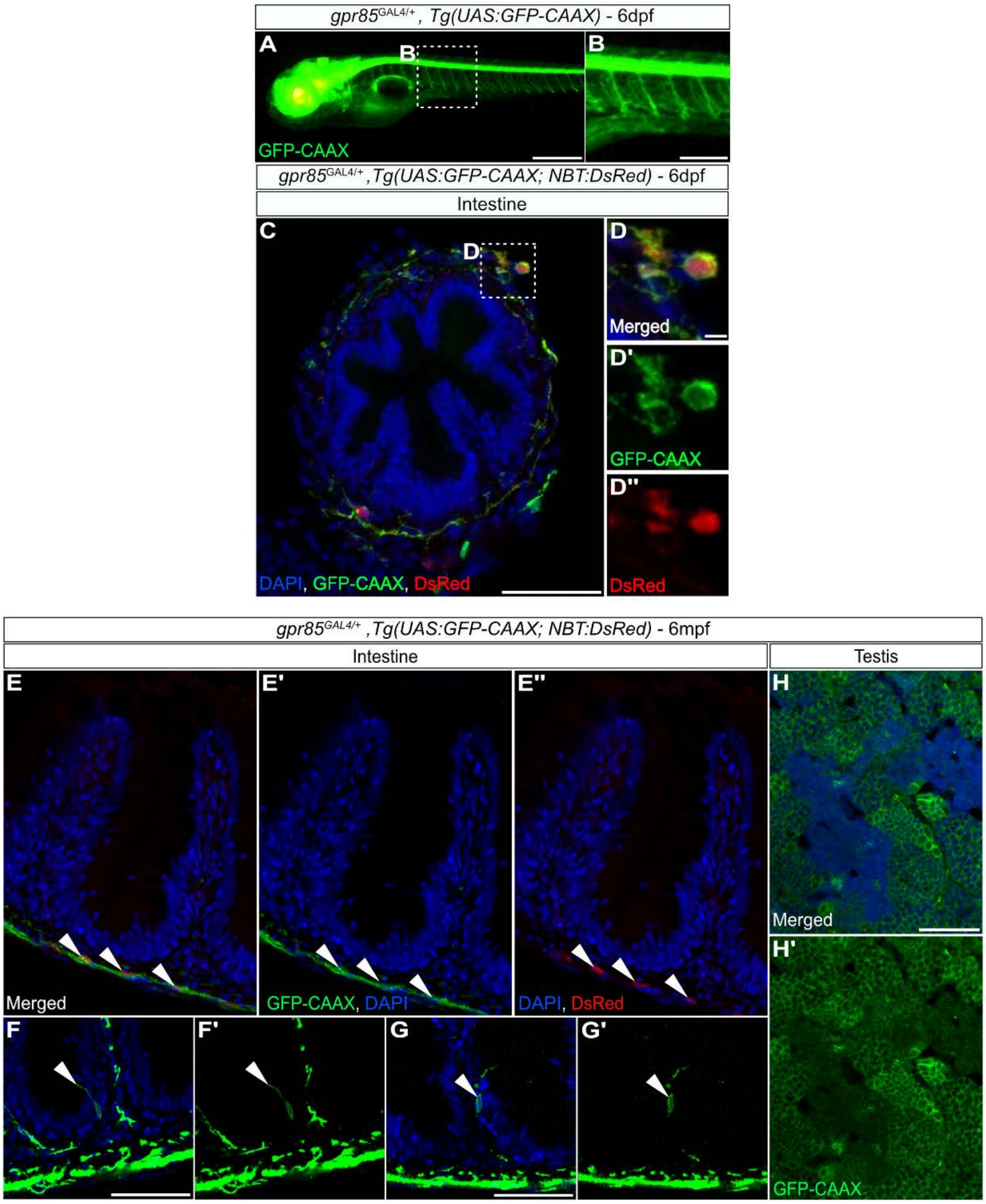


### Appendix Figure S2

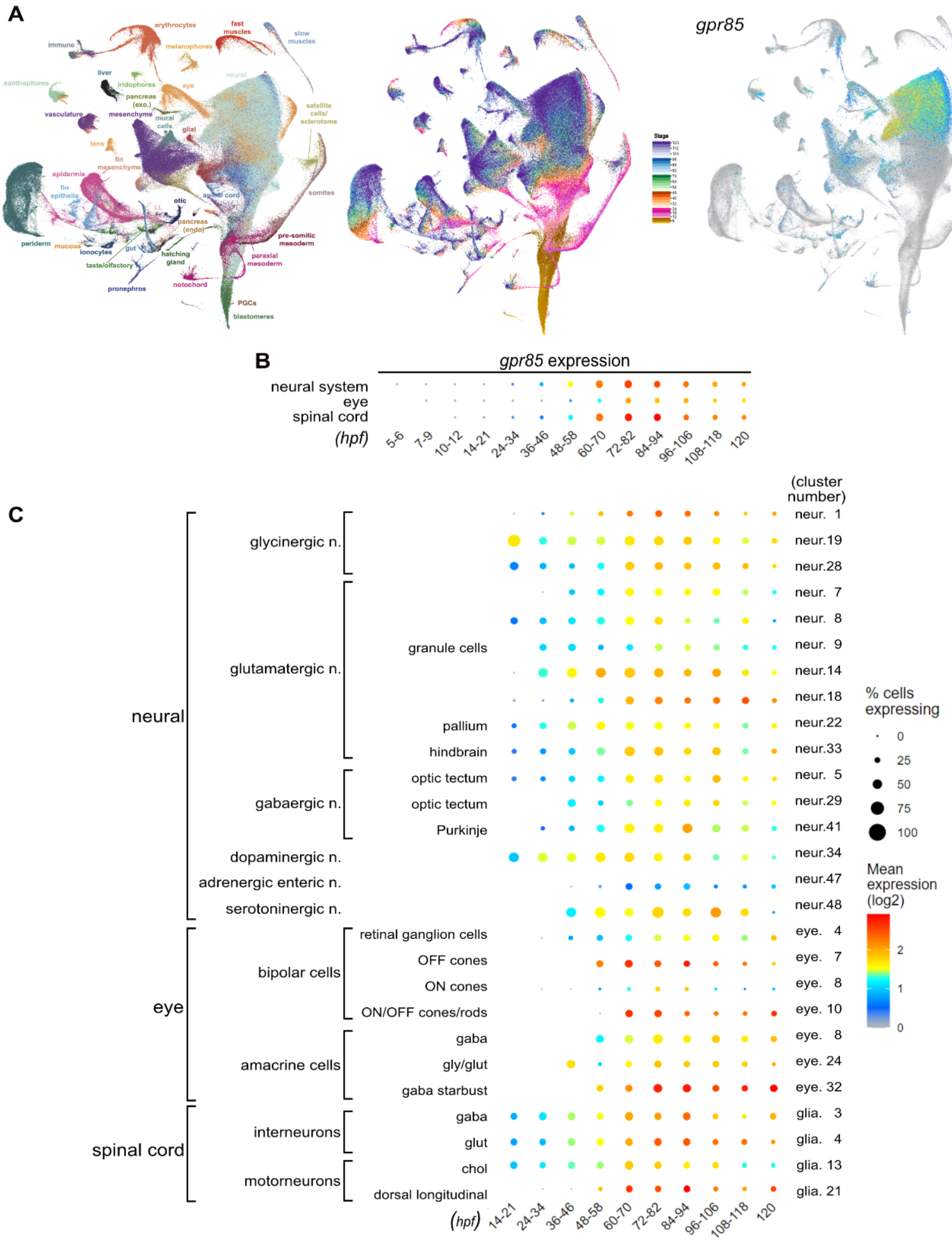


**Appendix Figure S3**

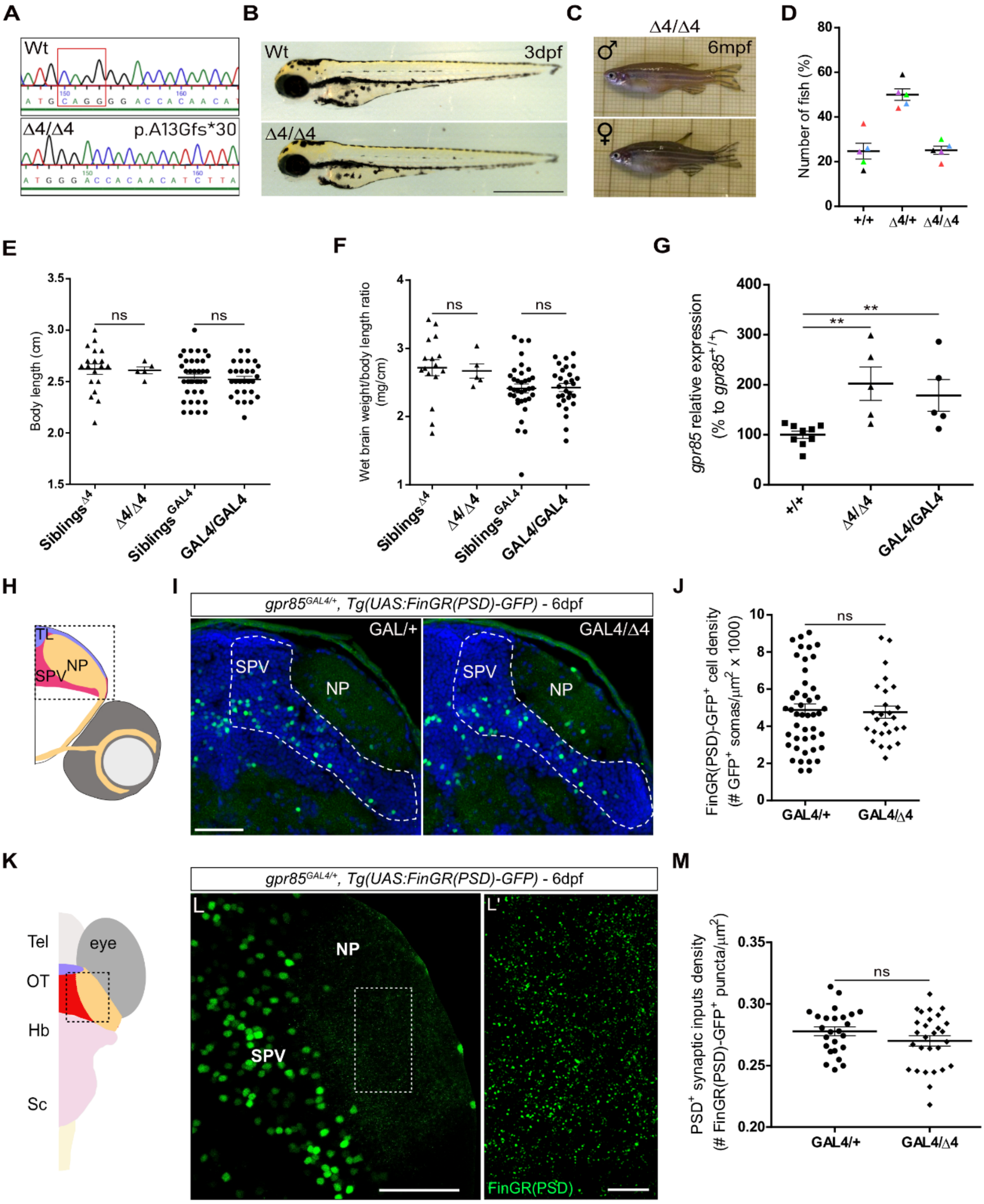


**Appendix Figure S4**

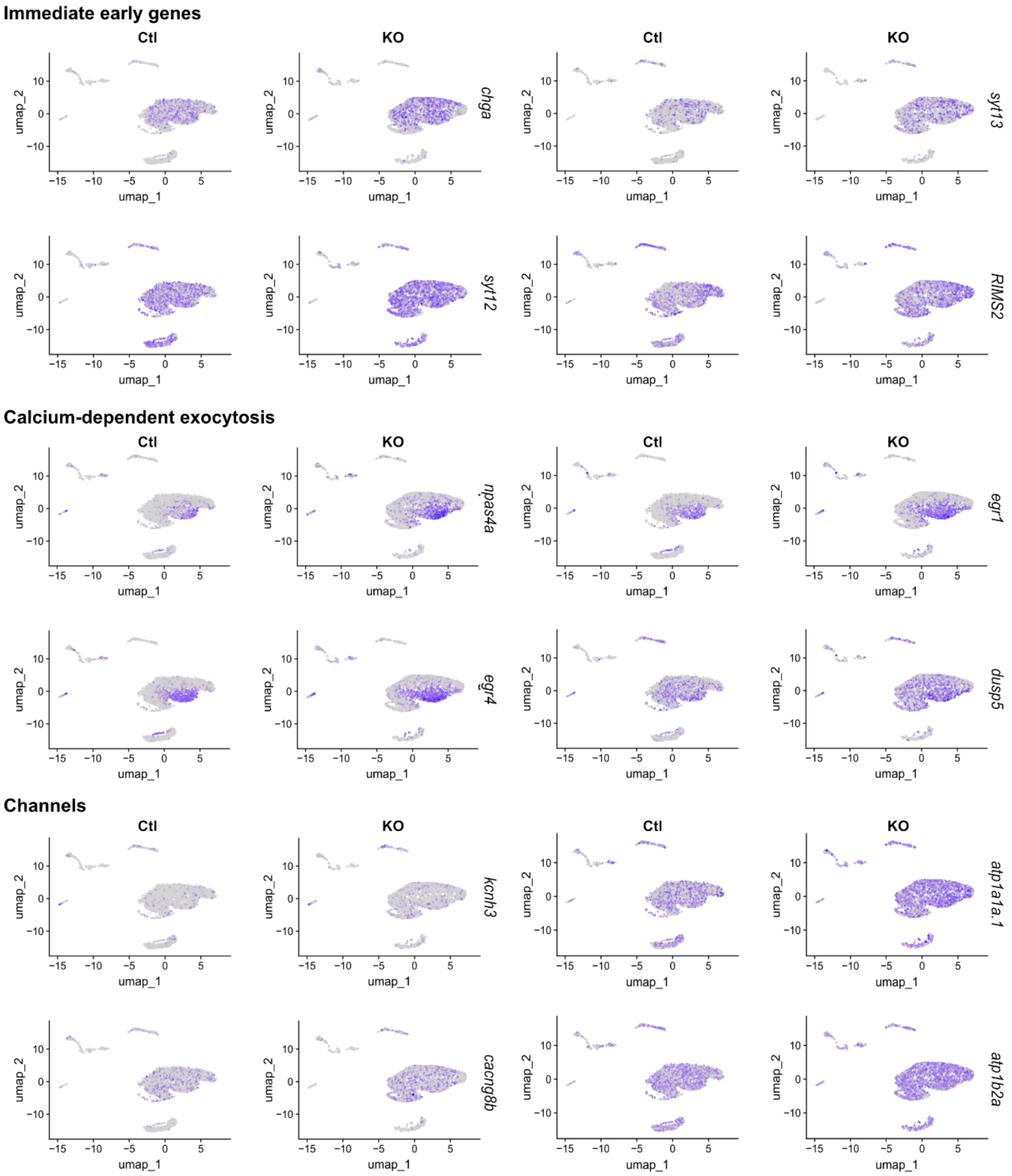


Appendix Figure S5

| Gene name | FC | adj_p_val |
| --- | --- | --- |
| IEG | | |
| *junba* | 3,89 | 1,387E-19 |
| *npas4a* | 3,57 | 4,675E-52 |
| *fosab* | 2,56 | 3,506E-12 |
| *egr1* | 2,45 | 2,675E-37 |
| *egr4* | 2,42 | 8,418E-39 |
| *junbb* | 2,32 | 1,393E-23 |
| *gadd45ba* | 2,21 | 3,104E-20 |
| *ier2b* | 2,14 | 3,978E-18 |
| *ier2a* | 1,83 | 3,98E-15 |
| *dusp5* | 1,69 | 2,844E-27 |
| EXOCYTOSIS | | |
| *sytl5* | 2,60 | 6,098E-21 |
| *chga* | 1,94 | 7,318E-53 |
| *syt14b* | 1,79 | 1,675E-26 |
| *scamp5a* | 1,72 | 7,662E-18 |
| *syt13* | 1,68 | 8,646E-32 |
| *cplx2.1* | 1,64 | 1,401E-19 |
| *cadpsa* | 1,55 | 2,068E-16 |
| *syt7b* | 1,44 | 2,411E-10 |
| *RIMS2* | 1,43 | 2,3E-27 |
| *syn2b* | 1,41 | 4,149E-08 |
| *syt12* | 1,39 | 5,378E-38 |
| CHANNELS | | |
| *atp1a3b* | 18,31 | 2,608E-63 |
| *clcn2a* | 3,11 | 2,224E-14 |
| *cacna1ba* | 2,45 | 2,126E-09 |
| *atp1b1b* | 2,10 | 3,521E-30 |
| *kcnh3* | 2,00 | 2,523E-23 |
| *atp1a1a.1* | 1,84 | 2,9E-101 |
| *cacng8b* | 1,59 | 9,476E-19 |
| *atp1b2a* | 1,58 | 5,123E-80 |

**Appendix Table S1**

### Appendix Figures Legends

#### Appendix Figure S1 - Whole-mount *in situ* hybridization and a newly generated knock-in reporter line highlight the predominant expression of gpr85 in the CNS and retina of zebrafish embryos.

**A,** Lateral views of whole-mount in situ hybridization of wild-type zebrafish embryos stained with antisense or sense *gpr85* probes at 1, 2, 3 and 6 dpf. BM purple was used for detection. At 1 dpf, *gpr85* expression is observed in the olfactory bulb, pallium, subpallium region (black arrowhead), and in the hypothalamic region of the telencephalon (white arrowhead). Scale bars: 500 µm. **B,** Magnified view of in situ hybridized eyes from 3dpf wild-type zebrafish embryos. BM purple was used for detection, and gpr85 transcripts were observed in the retina. Scale bars: 50 µm. **C,** Schematic diagram of the CRISPR/cas9 homology repair (HR) approach used to generate the *gpr85*$^{GAL4}$ knock-in reporter line. The sgRNA target sequence is shown in red, the arrow indicates the Cas9 cutting site and the protospacer adjacent motif (PAM) is underlined. The HR-donor vector contains 24 pb left (LH) and right (RH) homology arms surrounding the GAL4 open reading frame. **D,** Insertion of the GAL4 cassette was confirmed by PCR genotyping. The *gpr85*$^{+}$ allele produces a 300 bp amplicon, while the *gpr85*$^{GAL4}$ allele produces a 550 bp amplicon. **E-H,** Maximum projection confocal images of retinal coronal sections from *gpr85*$^{GAL4/+}$, *Tg(UAS:GFP-CAAX)* fish at 3dpf (E) and 6 dpf (G). Boxed regions are enlarged in panels F and H. Single channels show staining with anti-GFP antibody (green; F', H'), and DAPI (blue; F'', H''). GFP-CAAX$^{+}$ cells are visible in the inner nuclear layer (INL), and ganglion cell layer (GCL) but not in the photoreceptor layer (ONL). Scale bars: (E, G) 50 µm; (F, H) 25 µm. **I-J,** Maximum projection confocal images of coronal sections of the spinal cord from *gpr85*$^{GAL4/+}$, *Tg(UAS:GFP-CAAX)* zebrafish larvae at 3 dpf (I) and 6 dpf (J) stained with anti-GFP antibody (green; I'-J'), and DAPI (blue; I''-J''). GFP-CAAX$^{+}$ somas are observed in the ventral and dorsal parts. Scale bars: 50 µm. Data information: dpf, days post-fertilization.

**Appendix Figure S2 - The Gpr85 reporter line reveals gpr85-expressing neurons in the developing and adult intestine and in the testis.**

**A,** Lateral view of a live 6 dpf *gpr85*$^{GAL4/+}$ *,Tg(UAS:GFP-CAAX)* larva. The box indicates the region depicted in panel B. Scale bar: 500 µm. **B,** Higher magnification of the boxed area in A focusing on the intestinal tract. The signal intensity was enhanced, highlighting GFP-CAAX$^{+}$ cells in the gut wall (white arrows). Scale bar: 250 µm. **C,** Representative maximum projection confocal image of an intestinal coronal section from a 6 dpf *gpr85*$^{GAL4/+}$ *,Tg(UAS:GFP-CAAX, NBT:DsRed)* larva, stained with anti-GFP (green), anti-DsRed (red), and DAPI (blue). The box indicates the region enlarged in panel D. Scale bars: 50 µm. **D,** Higher magnification of the region boxed in panel C, showing a GFP-CAAX$^{+}$/DsRed$^{+}$ neuron. Scale bars: 10 µm. **E-G,** Maximum projection confocal images of intestinal coronal sections from adult *gpr85*$^{GAL4/+}$*, Tg(UAS:GFP-CAAX; NBT:DsRed)* zebrafish stained with anti-GFP (green), anti-DsRed (red), and DAPI (blue). White arrowheads indicate GFP-CAAX$^{+}$/DsRed$^{+}$ myenteric neurons (E-E") and GFP-CAAX$^{low}$ cells within the villi (F-G'). Scale bars: 50 µm. **H,** Representative maximum projection confocal image of testis sections from adult *gpr85*$^{GAL4/+}$*, Tg(UAS:GFP-CAAX)* zebrafish stained with anti-GFP (green) and DAPI (blue). Scale bar: 50 µm.

**Appendix Figure S3 - *Gpr85* is expressed during zebrafish development in brain, retina and spinal cord neurons.**

**A,** UMAP projections of the single-cell RNAseq time course from the Daniocell online resource (Farrell *et al*, 2018; Sur *et al*, 2023; Data ref: Sur *et al*, 2023). From left to right, UMAPs display the annotation of identified clusters, developmental stages and *gpr85* relative expression, respectively. **B,** The dot plot highlights the relative *gpr85* expression across developmental stages from 5-6 to 120 hpf, focusing on the neural, eye and spinal cord clusters. **C,** Dot plots of neuronal, retinal, and spinal cord subclusters showing *gpr85* the relative expression pattern from 14-21 to 120 hpf stages. All dot plots are accessible at https://daniocell.nichd.nih.gov/gene/G/gpr85/gpr85.html. Data information: hpf, hours post fertilization, scales (% cells expressing, Mean expression) shown in C also apply to B.

**Appendix Figure S4 - Zebrafish *gpr85* loss-of-function models are phenotypically normal, and cell and excitatory synaptic input densities of *Gpr85*-deficient neurons are unaffected through development.**

**A,** Genomic sequence of wild-type (Wt) and *gpr85* deletion mutant (Δ4/Δ4) zebrafish. The mutant *gpr85* sequence, generated via CRISPR/Cas9, results in a 4 base pair deletion (boxed) in exon 2, introducing a premature stop codon 17 amino acids later (p.Ala13Glyfs*30). **B,** No phenotypic differences were observed between 3 dpf Wt or Δ4/Δ4 larvae. Scale bar: 500 µm. **C,** Adult *gpr85*$^{\Delta4/\Delta4}$ zebrafish show no macroscopic defects. **D,** Genotypic analysis of adult fish from heterozygous intercrosses. A normal mendelian inheritance ratio is observed. Each dot color represents an independent cross (n = 5, at least 25 fish genotyped per cross, means: +/+, 24.7%; +/Δ4, 50%; Δ4/Δ4, 25.3%). **E,** Body length measurements of adult zebrafish with *gpr85*$^{\Delta4/\Delta4}$ (n = 5) and *gpr85*$^{GAL4/GAL4}$ (n = 28) loss-of-function mutations, compared to their respective control siblings (n = 18 for Δ4 siblings; n = 37 for GAL4 siblings). Statistical analysis: +/+ vs Δ4/Δ4: ns, p=0,68; GAL4/+ vs GAL4/GAL4: ns, p=0.59, Mann-Whitney. **F,** Wet brain weight normalized to body length for adult zebrafish with *gpr85*$^{\Delta4/\Delta4}$ and *gpr85*$^{GAL4/GAL4}$ loss-of-function mutations, compared to their respective control siblings (+/+ vs Δ4/Δ4: ns, p = 0.41; GAL4/+ vs GAL4/GAL4: ns, p = 0.91, Mann-Whitney test). **G,** *gpr85* transcript levels measured by RT-QPCR in *gpr85*$^{+/+}$ (n = 9), *gpr85*$^{\Delta4/\Delta4}$ (n = 5) and *gpr85*$^{GAL4/GAL4}$ (n = 5) 6 dpf larvae (+/+ vs Δ4/Δ4: **, p = 0.002; +/+ vs GAL4/GAL4: **, p = 0.007, Mann-Whitney test). **H,** Schematic coronal view of a 6 dpf zebrafish larva brain. The box indicates the optic tectum region imaged in I. **I,** Maximum projection confocal images of optic tectum coronal sections from 6 dpf *gpr85*$^{GAL4/+}$, *Tg(UAS:FinGR(PSD)-GFP)* larvae immunostained with anti-GFP (green) and DAPI (blue). Scale bar: 50 µm. **J,** Quantification of PSD$^{+}$ *gpr85*-expressing neuronal soma in the optic tectum SPV at 6 dpf. *gpr85*$^{GAL4/+}$ controls (n = 45) are compared with *gpr85*$^{GAL4/\Delta4}$ fish (n = 26), (*gpr85*$^{GAL4/+}$ vs *gpr85*$^{GAL4/Δ4}$, ns: not significant, p = 0.8, unpaired t-test). **K,** Schematic dorsal view of a 6 dpf zebrafish larva brain. The box indicates the optic tectum region imaged in L. **L,** Live confocal imaging of the optic tectum area from a 6 dpf *gpr85*$^{GAL4/+}$, *Tg(UAS:FinGR(PSD)-GFP)* larva (dorsal view). The boxed region is enlarged in L', showing PSD$^{+}$ excitatory synapses from *gpr85*-expressing neurons. Scale bars: L, 50 µm, L', 10 µm. **M,** Quantification of PSD$^{+}$ excitatory synapses belonging to *gpr85*-expressing neurons in the optic tectum neuropil at 6 dpf. *gpr85*$^{GAL4/+}$ controls (n = 25) are compared with *gpr85*$^{GAL4/GAL4}$ fish, as *gpr85*-deficient model (n = 28), (*gpr85*$^{GAL4/+}$ vs *gpr85*$^{GAL4/Δ4}$, ns: not significant, p = 0.169, unpaired t-test). Data information: SPV: stratum paraventriculare, NP: neuropil. Data are presented as mean ± SEM.

### Appendix Figure S5 - scRNAseq supplementary UMAPs Gpr85-Ctl vs Gpr85-KO

**A,** UMAPs of the Ctl and KO cells displaying the expression of DEG mentioned in Figure 5D. These plots show that the upregulation of IEGs is mostly found in cluster 2, while other genes related to neuronal activity exhibit broader upregulation across the GC populations.

### Appendix Tables Legends

### Appendix Table S1 - List of the genes related to neuronal activity up-regulated in *gpr85*-KO granule cells.

Data information: FC: Fold change; IEC: immediate early genes; adj_p_val: adjusted p-value
