## Supplementary material for "Synaptic Gpr85 influences cerebellar granule cells electrical properties and light-induced behavior": Reagents and tools table1

**Reagents and Tools Table**

| **Reagent/Resource** | **Reference or Source** | **Identifier or Catalog Number** |
| --- | --- | --- |
| **Experimental Models** |  |  |
| *Tg(UAS:GFP-CAAX)*^m1230^ (danio rerio) | Fernandes *et al*, 2012 | Referred as *Tg(UAS:GFP-CAAX)* |
| *Tg(UAS:lifeact-eGFP)*^mu271^ (danio rerio) | Helker *et al*, 2013 | Referred as *Tg(UAS:lifeact-eGFP)* |
| *Tg(Xltubb:DsRed)*^zf148^ (danio rerio) | Peri and Nüsslein-Volhard, 2008 | Referred as *Tg(NBT:DsRed)* |
| *Tg(zcUAS:FinGR(PSD95)-GFP-ZFC(CCR5TC)-KRAB(A)) (danio rerio)* | This study | Referred as *Tg(UAS:FinGR(PSD)-GFP)* |
| *Tg(ziUAS:FinGR(GPHN)-mCherry-ZFI(IL2RGTC)-KRAB(A)) (danio rerio)* | This study | Referred as *Tg(UAS:FinGR(GPHN)-mCherry)* |
| *Tg(UAS:zGpr85-eGFP) (danio rerio)* | This study |  |
| *Tg(UAS:zGpr85-mCherry) (danio rerio)* | This study |  |
| *Gpr85*^GAL4^ (danio rerio) | This study |  |
| *Gpr85^Δ^*^4^ (danio rerio) | This study |  |
| **Recombinant DNA** |  |  |
| *tol2*-UAS-MCS-EGFP | Genecust, Puc57 customed |  |
| *tol2*-UAS:zGpr85-eGFP | This study |  |
| *tol2*-UAS:zGpr85-mCherry | This study |  |
| pTol2-zcUAS:PSD95.FinGR-GFP-ZFC(CCR5TC)-KRAB(A) | Son *et al*, 2016 | Addgene plasmid #72638 |
| pTol2-ziUAS:GPHN.FinGR-mCherry-ZFI(IL2RGTC)-KRAB(A) | Son *et al*, 2016 | Addgene plasmid #72639 |
| **Antibodies** |  |  |
| chicken anti-GFP polyclonal | Abcam | ab13970, 1:1000 |
| mouse anti-mCherry monoclonal | Takara | 632543, 1:1000 |
| mouse anti-HuC/D monoclonal | Invitrogen | A-21271, 1:250 |
| mouse anti-PCNA monoclonal | Dako | M0879, 1:250 |
| rabbit anti-DsRed polyclonal | Takara | 632496, 1:1000, 1:50 |
| rabbit anti-Ribeye-A polyclonal | Zenisek’s lab | s4561-2, 1:1000, 1:50 |
| goat Alexa Fluor 488-conjugated anti-chicken IgG antibody | Abcam | ab150169, 1:500 |
| donkey Alexa Fluor 594-conjugated anti-rabbit IgG | Abcam | ab150076, 1:500, 1:100 |
| donkey Alexa Fluor 594-conjugated anti-mouse IgG | Abcam | ab150108, 1:500 |
| **Oligonucleotides and other sequence-based reagents** |  |  |
| *gpr85*-targeting sgRNA1 | This study | 5’-GGCGAACTACAGCCATGCAG -3’ |
| *gpr85*-targeting sgRNA2 | This study | 5’-AGTTCGCCATAGATGGAGAA-3’ |
| *UgRNA* | Wierson *et al*, 2020; Jordan M *et al*, 2021 | 5’-GGGAGGCGUUCGGGCCACAGCGG -3’ |
| GAL4 customed donor vector | This study | Left homology arm (5’-AGATCATGATCCTTGGCTAAAGCTTTAAGCGTTCTCTATGATCCCTTC-3’)/ Right homology arm (5’-TCCATCTATGGCGAACTACAGCCATGCAGGGGACCACAACATCTTACA-3’) |
| *gpr85*-Forward primer genotyping | This study | 5′-GAGACAAAGGAACAAAGGATGC-3’ |
| *gpr85*-Reverse primer genotyping | This study | 5′-CAGGATGGATATCAGGAGGTTT-3’ |
| GAL4-Reverse primer genotyping | This study | 5′-TGCTGTCTCAATGTTAGAGGCATATC-3’ |
| WISH *gpr85*-asFw primer | This study | 5’- TCAAAGACAAGAGCCTTCATCG-3’ |
| WISH *gpr85*-asRv primer | This study | 5’- GGATCCATTAACCCTCACTAAAGGGAACCTGCTTATCCGCTTTTCAGTT-3’ |
| WISH *gpr85*-sFw primer | This study | 5’- GGATCCATTAACCCTCACTAAAGGGAATCAAAGACAAGAGCCTTCATCG-3’ |
| WISH *gpr85*-sRv primer | This study | 5’- CCTGCTTATCCGCTTTTCAGTT-3’. |
| *qPCR gpr85*-Fw primer | This study | 5′-gcatcttctccaaccgagag-3’ |
| *qPCR gpr85*-Rv primer | This study | 5′-aaaacaaacgccacagaacc-3’ |
| *qPCR rpl13*-Fw primer | This study | 5′-TCTGGAGGACTGTAAGAGGTATGC-3’ |
| *qPCR rpl13*-Rv primer | This study | 5′- AGACGCACAATCTTGAGAGCAG-3’ |
| *qPCR lms12b*-Fw primer | This study | 5′-AGCCATGTCTCTTGCCTCAC-3’ |
| *qPCR lms12b*-Rv primer | This study | 5′-ATGACGTCACTGAGGTTGGG-3’ |
| Oligo(dT) primer | Thermo Fisher Scientific | 10304690 |
| **Chemicals, Enzymes and other reagents** |  |  |
| BM-purple | Roche | 11442074001 |
| TRIzol reagent | Qiagen | 79306 |
| PFA | Sigma-Aldrich | 158127 |
| Sucrose | Millipore | 1076511000 |
| THF | Sigma-Aldrich | 186562 |
| Triethylamine | Sigma-Aldrich | T0886 |
| Nycodenz | Accurate Chemical & Scientific | AN1002423 |
| Glutaraldehyde | Sigma-Aldrich | 340855 |
| Papain | Sigma-Aldrich | p4762 |
| DPBS | Gibco | 14190144 |
| FBS | Gibco | A5256701 |
| Normal goat serum | abcam | Ab7481 |
| Normal donkey serum | abcam | Ab7475 |
| Sytox Red | Thermofisher scientific | S34859 |
| Chromium Single cell 3’ kit v3 | 10x genomics | PN-1000268 |
| Cas9 | PNA Bio | CP01-50 |
| *Xho*I | New England Biolabs | R0146S |
| *Bbv*CI | New England Biolabs | R0601S |
| *Sal*I | New England Biolabs | R0138S |
| RNAeasy Mini Kit | Qiagen | 74104 |
| TURBO DNA-*free* kit | Thermo Fisher Scientific | AM1907 |
| Super Script II reverse transcriptase | Thermo Fisher Scientific | 18064014 |
| Tissue freezing medium | Leica | 14020108926 |
| Agar 100 Resin Kit | Agar scientific | AGR1031 |
| **Software** |  |  |
| Seurat package in R | Stuart *et al*, 2019 |  |
| SCTransform | Hafemeister and Satija, 2019 |  |
| Biorender | https://www.biorender.com/ |  |
| Rstudio |  |  |
| GraphPad Prism | https://www.graphpad.com |  |
| Fiji | https://imagej.net/software/fiji |  |
| iTEM | Olympus |  |
| LAS V4.6.2 | Leica |  |
| Imaris | Oxford instruments |  |
| NIS-Elements AR | Nikon |  |
| PatchMaster | HEKA |  |
| Zen | Zeiss |  |
| IgorPro 6.3 | WaveMetrics |  |
| Axograph X | Axon Instruments Inc. |  |
| Zebralab | Viewpoint |  |
| **Other** |  |  |
| Cryostat | Leica | CM3050 S |
| Precision balance | Sartorius Entris II | BCE224I-1S |
| Ultramicrotome | Leica EM | UC7 |
| Cryo-chamber | Leica EM | FC6 |
| Transmission electron microscope | FEI, Thermo Fisher Scientific | Tecnai 10 100kV |
| Fluorescent stereomicroscope | Leica | M165FC |
| Digital camera | Leica | DFC7000T |
| Inverted confocal microscope | Zeiss | LSM 780 |
| Confocal microscope | Nikon | AX/R |
| 26 G needle | B.Braun | Omnifix 100 Duo |
| Vibratome | Leica | R VT 1000 S |
| Axioskop microscope | Zeiss | 2FS Plus |
| Infrared CCD camera | Hamamatsu Photonics KK | XST70CE |
| Electroluminescent diode | Cairn Research Lda | OptoLED R |
| Patch clamp amplifier | HEKA | EPC-10 |
| 48 well-plates | VWR | 77589 |
| 40 µm nylon mesh | VWR | 76327-098 |
| Carbon-formvar 100 mesh copper grids | Electron Microscopy Sciences | 501903323 |
| Borosilicate-glass patch electrodes | GmbH |  |
| Chromium controller-10X Genomics | 10X Genomics |  |
| NovaSeq 6000 | Illumina |  |
| FACS ARIA II | BD Biosciences |  |
| CFX96 Real-Time System | Bio-rad, CA |  |
| Zebrabox | Viewpoint |  |
| MegaView 14 bits camera | Olympus |  |
